# Precision mapping and molecular contextualization of surgical outcome epicenters in temporal lobe epilepsy

**DOI:** 10.64898/2026.03.06.710165

**Authors:** Fatemeh Fadaie, Ke Xie, Jack Lam, Thaera Arafat, Ella Sahlas, Judy Chen, Jessica Royer, Ian Goodall-Halliwell, Rui Ding, Marlo Naish, Raúl Rodríguez-Cruces, Jiajie Mo, Jeffery Hall, Yifei Weng, Sara Larivière, Sami Obaid, Aristides Hadjinicolaou, Alexander G. Weil, Raluca Pana, Zhiqiang Zhang, Andrea Bernasconi, Neda Bernasconi, Boris Bernhardt

**Author notes:** **Correspondence to:** Fatemeh Fadaie -, Boris C. Bernhardt -, Multimodal Imaging and Connectome Analysis Laboratory, McConnell Brain Imaging Centre, Montreal Neurological Institute, McGill University, Montreal, Quebec, Canada.

## Abstract

Temporal lobe epilepsy is the most common drug-resistant epilepsy, with surgical resection offering the primary path to seizure freedom. Despite standardized approaches, a substantial proportion of patients experience seizure recurrence, and the neurobiological substrates underlying these divergent outcomes remain unclear. We applied an individualized normative modeling framework to multimodal preoperative MRI data in a group that subsequently underwent surgical resection, to characterize patient-specific structural deviations and identify disease epicenters. Patients who became seizure-free exhibited spatially coherent abnormalities localized to the hippocampus and ipsilateral association regions, anchored in agranular limbic territories and enriched for genes linked to calcium-dependent signaling. Non-seizure-free patients, on the other hand, showed a more heterogeneous and distributed pattern of deviations, consistent with a “temporal-plus” network organization, and broader neuromodulatory dysregulation. Crucially, overlap between resected tissue and network-defined epicenters was closely associated with seizure freedom, independent of total resection volume. These findings provide a multiscale framework for precision surgical planning, shifting the focus from standardized tissue removal to targeted disconnection of patient-specific pathological hubs.

## Introduction

Temporal lobe epilepsy (TLE) is widely considered as a prototypical surgically-amenable epilepsy, with a majority of patients showing seizure freedom and markedly improved quality of life after resection of the epileptogenic temporal lobe ^1,2^. However, approximately 30% of patients still continue to experience seizures after resection ^3,4^, often facing persistent cognitive, social, and safety challenges ^5^. Notably, despite innovations in diagnostic methods and surgical techniques, surgical success rates have not markedly improved over the last years ^6^. Previous studies have identified key contributors to seizure freedom, including the extent of resection of epileptogenic mesial temporal structures ^7–9^. More recent work has extended this view by implicating extra-temporal structural and functional networks, highlighting the influence of distributed alterations on postoperative seizure control ^10,11^. Despite extensive prior work demonstrating marked inter-patient variability in the neuroimaging signatures of TLE, both within the mesiotemporal lobe and across distributed cortical and subcortical systems ^12,13^, conventional group-level comparisons obscure patient-specific differences, limiting their prognostic value. At the hippocampal level, some patients show clear atrophy ^14–16^, whereas others display only subtle or no detectable structural abnormalities ^17–19^. Variability in neocortical and white matter systems has been studied through analyses of patient-specific patterns of cortical thinning and diffusion abnormalities, which affect mesiotemporal, temporo-frontal, or interhemispheric pathways to varying degrees ^12,13,20^. Normative modeling offers a solution by quantifying deviations from healthy populations to reveal patient-specific patterns across distributed brain systems, enabling individualized characterization and prognostic biomarker discovery, including prediction of surgical outcomes ^12,21–23^.

A natural extension of this approach is to ask whether individualized deviations converge on focal regions that serve as anchors of disease expression, so-called *disease epicenters*. The concept of disease epicenters has recently emerged as a framework to explain how focal pathology can give rise to widespread network abnormalities ^23–25^. Importantly, epicenters may differ in their cytoarchitecture and microstructural composition, reflecting regional variations in neuronal density, laminar organization, and local circuitry that could influence susceptibility to epileptic activity ^26–28^. In epilepsy, recent applications of epicenter mapping has revealed that focal lesions and network-level alterations often anchor to specific cortical or subcortical regions with characteristic connectivity profiles ^22,23,25,29^. Identifying these regions is clinically relevant in TLE, where surgical outcomes remain heterogeneous despite standardized procedures. By comparing epicenters between seizure-free (TLE-SF) and non-seizure-free (TLE-NSF) patients, we can test whether differences in their location or network embedding contribute to surgical failure, thereby moving toward more precise, biology-informed predictors of postoperative outcome.

A key unresolved issue is why particular temporal lobe regions repeatedly emerge as epileptic epicenters, and why their persistence can sustain seizure recurrence. The disease process in TLE is determined by diverse molecular and cellular factors, including disrupted synaptic milieu, excitatory–inhibitory balance, ion channel function, axonal guidance, and synaptic plasticity ^30,31^. These changes occur alongside characteristic neuropathological processes, such as neuronal degeneration, reactive gliosis, and maladaptive circuit reorganization ^32,33^. Together, these findings underscore the role of molecular pathways in shaping the selective vulnerability of temporal lobe networks. Importantly, different disease mechanisms (ranging from neuroinflammation and gliosis to structural remodeling and channelopathies) may variably influence seizure recurrence after surgery. Mapping transcriptomic signatures onto patient-specific epicenters thus offers a way to test whether distinct biological programs underlie surgical success versus relapse.

We analyzed a cohort of 102 patients with pharmaco-resistant TLE who underwent surgical resection after multimodal 3T MRI investigations and for whom at least 1-year postoperative outcome data were available. Patients were recruited from two independent surgical centres. Using a patient-specific normative modeling framework, we quantified deviations in cortical thickness (CT), fractional anisotropy (FA), and apparent diffusion coefficient (ADC) relative to healthy controls and derived individualized maps of structural pathology. From these maps, we identified epicenters as regions showing maximal coupling between local abnormalities and whole-brain network architecture in TLE-SF and TLE-NSF patients. These epicenters capture the topographically and functionally most salient loci of disease expression linking macrostructural network mechanisms to brain microstructure, molecular milieu, and surgical target definition. Accordingly, we examined their cytoarchitectural and transcriptomic context to uncover cellular and molecular pathways differentiating outcome groups and assessed their spatial overlap with surgical resections to elucidate anatomical determinants of surgical success and failure.

## Results

### Clinical and demographic characteristic

A total of 102 pharmaco-resistant TLE patients and 94 matched healthy controls were included from three independent cohorts (MICs, NOEL, Nanj) aggregated across two surgical centres (Montreal Neurological Institute and Hospital, Jinling Hospital Nanjing). Patients were classified as TLE-SF (n=77; Engel class I) or TLE-NSF (n=25; Engel classes II–IV) based on the surgical outcome, with a mean postoperative follow-up of 4.89±2.5 years. Histopathological evaluation revealed a significantly higher proportion of hippocampal cell loss with gliosis relative to isolated gliosis in TLE-SF patients compared to those with persistent seizures (66/16 vs. 11/9; χ²=4.35, P=0.03). No significant group differences were observed between TLE-SF and TLE-NSF patients in age at surgery, sex distribution, age at seizure onset, or seizure laterality (P>0.05). Although TLE-SF patients showed a trend toward longer disease duration, this difference did not reach statistical significance (P=0.10). Demographic and clinical characteristics for each cohort are summarized in **Table 1**.

**Table 1.** Demographic and clinical information.

| *Dataset (n)* | *Groups (n)* | *Age (years)* | *Gender (F/M)* | *Seizure focus (L/R)* | *Disease onset (years)* | *Disease duration (years)* | *Histopathology (HS/Gliosis)* | *Engel 1 (%)* | *Follow-up (years)* |
| --- | --- | --- | --- | --- | --- | --- | --- | --- | --- |
| *All (196)* | CON (94) | 34.38±10.42 | (45/49) | - | - | - | - | - | - |
|  | TLE (102) | 32.56±10.47 | (58/44) | (46/56) | 16.16±10.59 | 16.40±11.24 | (82/20) | 75% | 4.89±2.55 |
| *MICs (34)* | CON (23) | 45.52±9.07 | (11/12) | - | - | - | - | - | - |
|  | TLE (11) | 41.63±10.15 | (7/4) | (4/7) | 23.56±15.49 | 18.07±11.36 | (7/4) | 81% | 6.60±2.37 |
| *NOEL (91)* | CON (40) | 33.47±7.54 | (17/23) | - | - | - | - | - | - |
|  | TLE (51) | 35.07±10.06 | (29/22) | (25/26) | 15.86±10.69 | 19.24±12.55 | (35/16) | 78% | 5.76±2.33 |
| *Nanj (71)* | CON (31) | 27.29±7.37 | (17/14) | - | - | - | - | - | - |
|  | TLE (40) | 26.87± 7.93 | (22/18) | (17/23) | 14.55±8.15 | 12.32±7.96 | (40/0) | 70% | 3.31±2.01 |

### Normative modeling analysis (Individualized *w*-score maps)

To capture heterogeneous structural alterations at the level of individual patients, we used a normative modeling framework on complementary markers of tissue integrity (*i.e.,* CT, FA, and ADC). Healthy control data were used to establish normative distributions, from which we derived age- and sex-adjusted w-scores for each region in each patient ^22,23^. Regions showing extreme deviations (|w|≥1.96) were identified, producing individualized structural abnormality maps that enabled robust comparison of deviation patterns across TLE subgroups (**Fig. 1a**). Compared to healthy controls, both TLE-SF and TLE-NSF groups exhibited extensive yet variable alterations in CT, FA, and ADC (**Fig. 1b**). Across the cortex, the median proportion of extreme deviations in CT, FA, and ADC in TLE-SF was 6.68% (range=1.33–40.9%), 9.35% (0–50%), and 9.62% (0.26–77.80%), respectively. In TLE-NSF, the corresponding values were 6.95% (range=2.94–16.84%), 11.22% (4.54–39.83%), and 10.69% (1.06–61.22%). Stratification by cytoarchitectural maps revealed that both groups showed hippocampal atrophy, FA decrease in association regions, and ADC increase in the hippocampus and secondary association cortex. TLE-NSF, however, showed additional cortical atrophy in the primary sensory cortex ipsilaterally and motor cortex contralaterally. On fine-grained hippocampal analysis, TLE-SF patients showed a predominantly anterior pattern of hippocampal atrophy, whereas TLE-NSF patients exhibited greater atrophy posteriorly. FA showed mild bilateral decreases in both groups. ADC increases were more pronounced ipsilaterally in both groups, with an additional subtle contralateral increase in TLE-NSF (**Fig. S1a**). Moreover, we observed that 16% and 12% of TLE-NSF patients showed ipsilateral amygdalar and hippocampal hypertrophy, respectively (**Fig. S2**), while no such hypertrophy was identified in TLE-SF.

**Figure 1.**
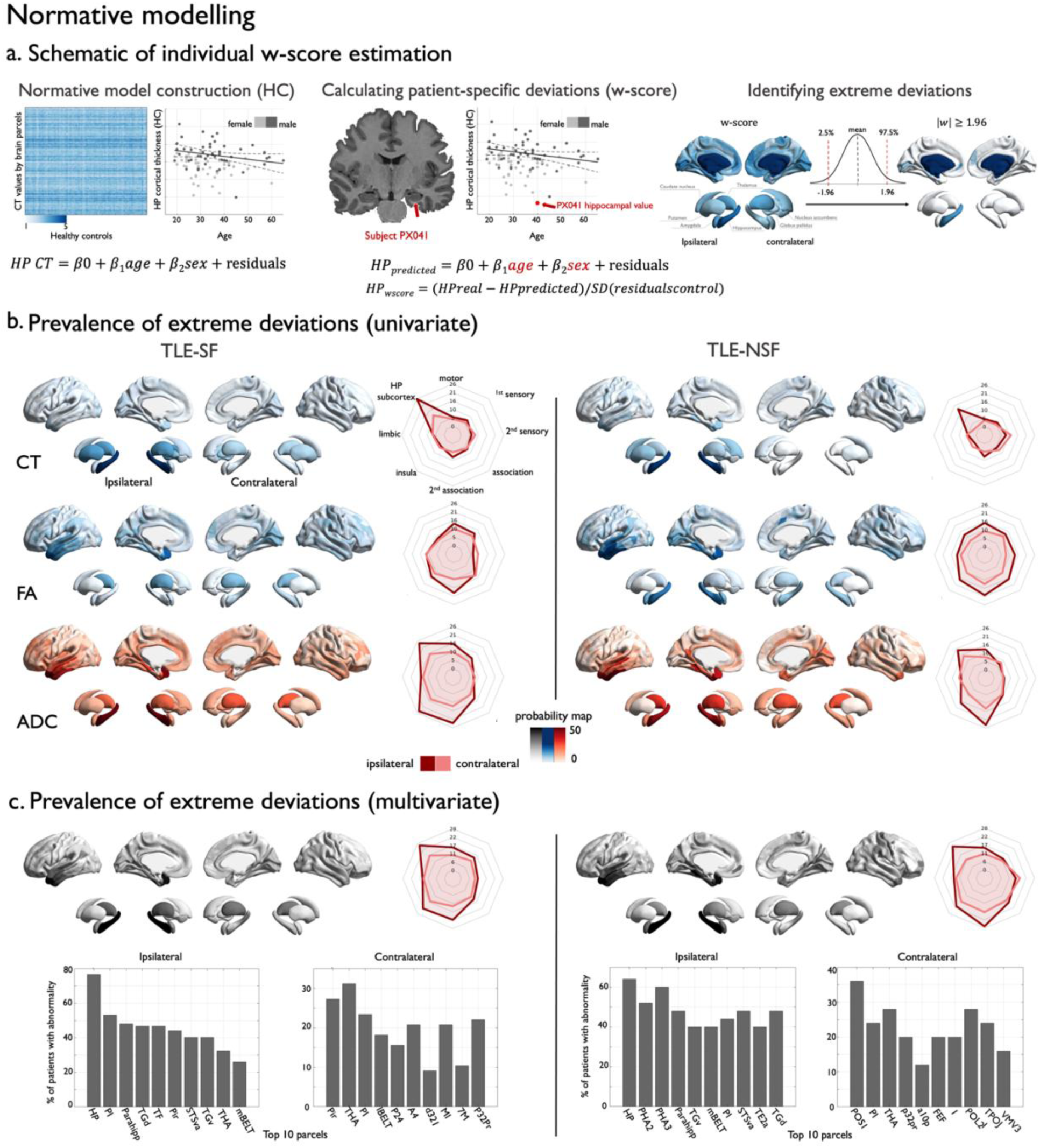
Normative modeling. **(a)** The left panel shows normative model generation in healthy controls, where age-and sex-related variations in hippocampal volume are modeled using linear regression, yielding beta maps for intercept (β0), age (β1), sex (β2), and standard deviation of residuals. The middle panel illustrates deviation estimation in a single patient: predicted hippocampal volume is calculated as β0 + β1×age + β2×sex, and w-scores are defined as the normalized deviation from predicted values. The MRI view highlights lower left hippocampal volume (*red flash*), and the scatter plot shows this patient’s value relative to healthy controls. Right panel shows thresholding of w-scores to identify extreme deviations (|w|≥1.96, corresponding to the upper and lower 2.5% of the normative distribution). **(b)** Proportion of TLE patients showing extreme deviations (|w|≥1.96) in cortical thickness (CT), fractional anisotropy (FA) and apparent diffusion coefficient (ADC) displayed separately for seizure-free (TLE-SF, *left*) and non–seizure-free (TLE-NSF, *right*) groups. Blue indicates decreases and red indicates increases relative to healthy controls. Spider plots summarize the mean proportion of extreme deviations across cytoarchitectonic classes from the Von Economo and Koskinas atlas. **(c)** Proportion of extreme deviations in the multivariate w-score for TLE-SF (*left*) and TLE-NSF (*right*) groups. Spider plots summarize deviations across cytoarchitectonic classes, and the bottom bar plot shows the percentage of patients with abnormalities (|multivariate w|≥1.96) in the top 10% of affected regions, separately for ipsilateral and contralateral hemispheres. *Abbreviations*: HP, hippocampus; PHA2 and PHA3, parahippocampal areas 2 and 3; PI, posterior insula; Pir, piriform cortex; TGd, dorsal temporal gyrus; TGv, ventral temporal gyrus; TF, temporal fusiform cortex; STSva, ventral anterior superior temporal sulcus; TE2a, anterior temporal area TE2; mBElt, medial belt auditory cortex; THA, thalamus.

Because CT, FA, and ADC reflect complementary aspects of tissue integrity, we next evaluated their joint contribution using a multivariate Mahalanobis distance calculation to identify regions showing convergent deviations. Aggregating the three metrics using a multivariate Mahalanobis distance analysis revealed distinct spatial patterns of structural deviations across the cortex between the groups (TLE-SF median: 9.09% [0.53–63.6%]; TLE-NSF median: 12.03% [3.74–43.58%]; **Fig. 1c**). In the TLE-SF group, deviations were relatively homogeneous, with the hippocampus being the most frequently affected region (∼80%; **Fig.1c; bar plot**), followed by secondary association cortices (superior temporal, parieto-temporo-occipital junction) and the ipsilateral insular cortex (**Fig.1c; spider plot**). In contrast, the TLE-NSF group displayed a more heterogeneous and individually variable pattern. Hippocampal deviations were slightly less common (∼65%; **Fig.1c; bar plot**), with greater prevalence of abnormalities in secondary association cortices compared to hippocampus and additional involvement of contralateral sensory regions (**Fig.1c; spider plot**). Hippocampal analysis revealed frequent deviations on the ipsilateral side in both groups, with TLE-SF showing more common alterations in the anterior portion, whereas TLE-NSF exhibited marked deviations distributed throughout the hippocampus (**Fig. S1b**). In line with the findings of the normative model, a direct comparison between TLE-SF and TLE-NSF showed a trend toward a higher prevalence of deviations in the posterior parahippocampal gyrus in TLE-NSF, as well as a higher prevalence of contralateral deviations in this cohort (**Fig. S3)**. Stratifying by seizure-focus laterality, left and right TLE-SF showed similar patterns of extreme hippocampal deviation. In contrast, left TLE-NSF exhibited the greatest deviation in the ipsilateral insular cortex, whereas right TLE-NSF showed the most extreme deviation in the ipsilateral secondary associative cortex (**Fig. S4**). Stratifying by histopathology, TLE-SF patients in both the HS and gliosis groups showed abnormalities largely confined to the temporal lobe. In contrast, TLE-NSF patients exhibited abnormalities extending into association cortices, with even more pronounced contralateral involvement in the gliosis subgroup (**Fig. S5**). Finally, we confirmed the robustness of these findings across all three study sites using 1,000 spin permutation tests to account for spatial autocorrelation in the neocortex and shuffle tests for subcortical structures, yielding significant spatial similarity (r values = 0.29–0.78, P_spin_< 0.08/ P_spin_< 0.001; **Fig. S6**).

### Disease epicenter mapping

To explore whether distributed alterations in TLE emerge from specific network hubs, we asked how closely each brain region’s typical connectivity pattern aligns with the patient-specific pattern of deviations. Using structural and functional connectivity maps derived from healthy controls, we calculated an “epicenter strength” for each cortical and subcortical region, reflecting how well its normal connectivity matches the observed abnormality distribution (**Fig. 2a**). This approach highlights potential hub regions that may drive or reflect disease spread, providing potential targets for intervention. We then applied this framework to compare epicenter patterns between TLE-SF and TLE-NSF subgroups. In TLE-SF, concordant structural–functional epicenters were localized to ipsilateral mesial temporal and association regions, including the hippocampus (P_spin_<0.05; **Fig. 2b. 2c, left**). In TLE-NSF, a broadly similar ipsilateral temporal pattern was present (P_spin_<0.05; **Fig. 2b. 2c, right**). However, hippocampal involvement was confined to functional epicenter mapping and was not mirrored structurally. Direct group comparisons showed that TLE-SF patients showed a trend toward stronger epicenter values in the ipsilateral orbitofrontal region, whereas TLE-NSF patients exhibited stronger values in the contralateral precentral area (P<0.05). Functionally, epicenter values were stronger in the ipsilateral DMN network in TLE-SF, while TLE-NSF patients showed more widespread involvement, including bilateral fronto-parietal regions, occipital cortex, and posterior temporal lobe, indicating a broader spatial distribution (P_FDR_<0.05; **Fig. 2d**). Taken together, while both groups shared a common core of temporal lobe epicenters, TLE-SF showed stronger hippocampal concordance of structural and functional epicenters and predominantly ipsilateral epicenters. On the other hand, TLE-NSF showed hippocampal epicenters only at the level of function together with more widespread network reorganization. Finally, we confirmed the robustness of these findings across all three datasets using 1,000 spin permutation tests to account for spatial autocorrelation in the neocortex and shuffle tests for subcortical structures, demonstrating significant spatial similarity (r=0.31–0.73; P_spin_<0.1 to P_spin_<0.0001; **Fig. S7**). We further performed a replication analysis by generating epicenter maps using an independent healthy control dataset, which yielded highly concordant results (r=0.91–0.98; P_spin_<0.00001 to P_spin_<0.001).

**Figure 2.**
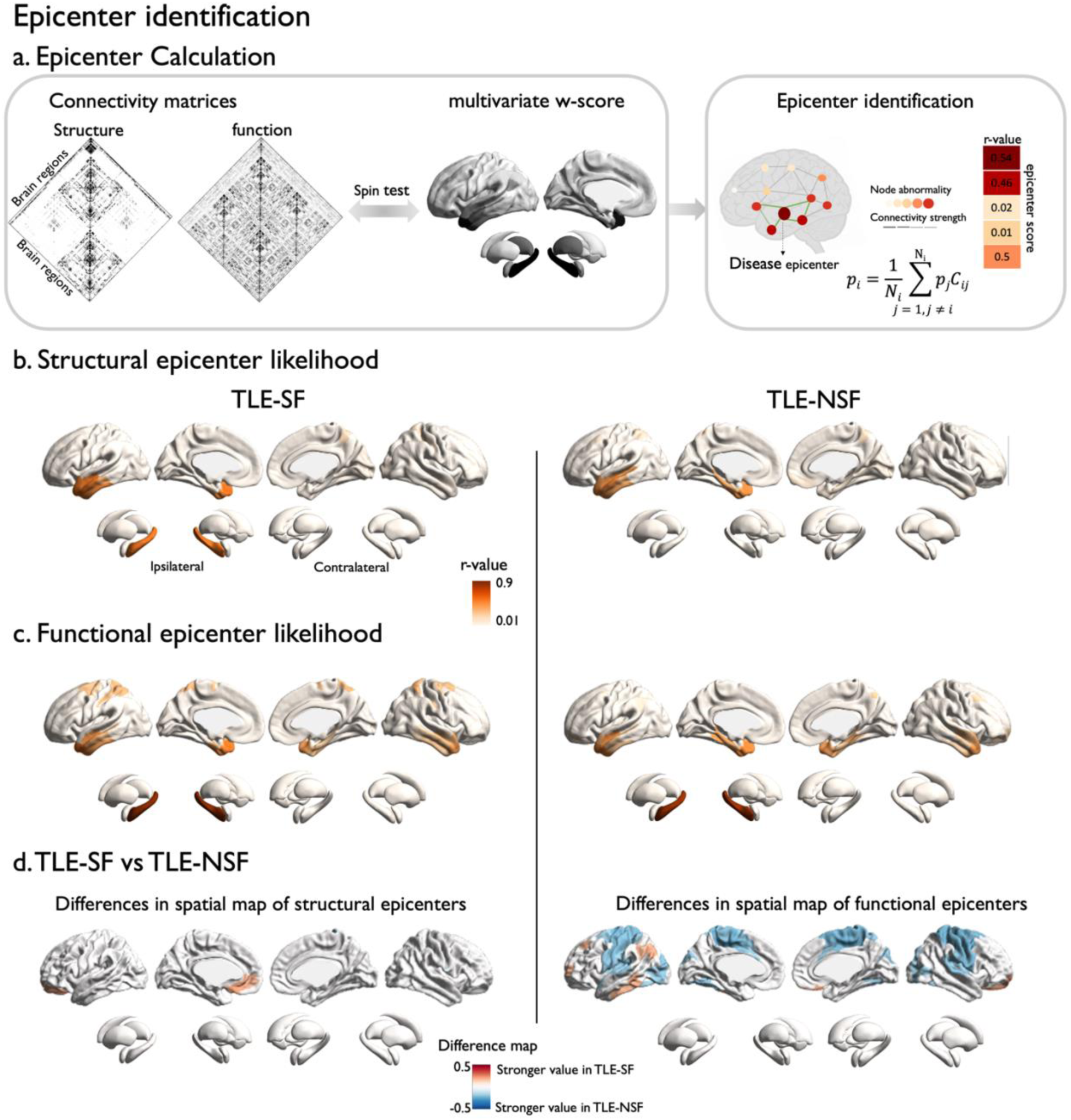
Epicenter identification. **(a)** Schematic of the epicenter calculation procedure. Left panel shows the matrix of structural and functional correlations with multivariate w-scores; right panel illustrates the workflow for identifying epicenters based on these maps. This approach captures a network-mediated effect, whereby regions connected to highly abnormal neighbors are more likely to show deviations, whereas regions connected to healthy neighbors are less likely to be affected. **(b-c)** Significant epicenters (TLE-SF: *left*; TLE-NSF: *right*) were visualized on cortical and subcortical surface templates for structure **(b)** and function **(c)**. **(d)** Differences in significant epicenters between TLE-SF and TLE-NSF were visualized on cortical and subcortical surfaces for structure *(left)* and function *(right)*. Red indicates regions where epicenter strength is greater in TLE-SF, and blue indicates regions where epicenter strength is greater in TLE-NSF.

### Association with cytoarchitecture maps

The pathophysiological network in TLE is thought to follow specific structural and microstructural pathways, suggesting that network epicenters may preferentially emerge in regions with distinctive cytoarchitectural properties. To test this, we related patient-derived epicenter maps to established cortical reference atlases: the Von Economo–Koskinas atlas capturing cell size, density, and thickness, and the BigBrain atlas characterizing laminar microstructure. Regional epicenter strength was quantified as the mean of structural and functional epicenter centrality for each parcel, computed separately for cortical and subcortical regions within the TLE-SF and TLE-NSF groups. Associations with cytoarchitectonic classes were assessed using spin permutation tests to account for spatial autocorrelation (**Fig. 3a**).

**Figure 3.**
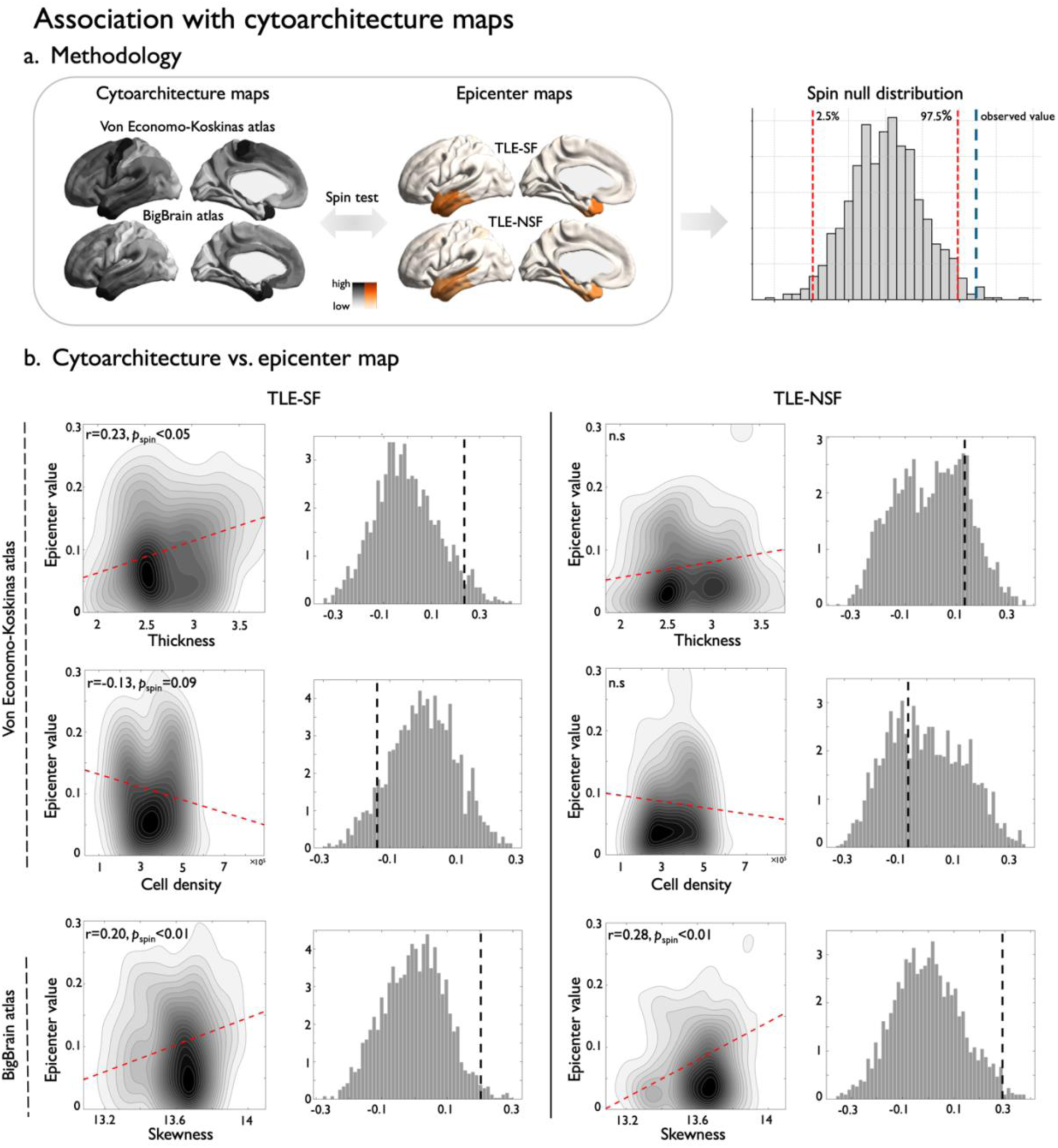
Association with cytoarchitecture maps. **(a)** Illustration of the spin test procedure showing correlations between mean structural-functional epicenter maps and cortical atlases while controlling for spatial autocorrelation. **(b)** Contour plots show correlations between cytoarchitectural features and epicenters for TLE-SF (*left*) and TLE-NSF (*right*). For each contour plot, the corresponding null distribution from the spin test is displayed to the right, with dashed black lines indicating observed significant correlations.

In TLE-SF, regional epicenter strength was higher in areas characterized by thicker cortex (r=0.23, P_spin_<0.05) according to the Von Economo–Koskinas atlas and there was a tendency for higher strength with larger cells as well (r=0.13, P_spin_=0.09) (**Fig. 3b**). Similarly, BigBrain-derived features showed stronger epicenters in regions with greater laminar skewness (r=0.20, P_spin_<0.01; **Fig. 3b**), reflecting a more differentiated and superficially weighted laminar structure. These associations are most prominent in limbic and paralimbic cortices, including mesial temporal and orbitofrontal regions, suggesting that structural–functional epicenters in TLE-SF preferentially emerge within evolutionarily older, integrative cortical territories. In contrast, the TLE-NSF group exhibited weaker and less spatially consistent associations, with significant increases observed only in laminar skewness (r=0.28, P_spin_<0.01; **Fig.3b**). This pattern may indicate that epicenters in TLE-NSF are more widely distributed across neocortical regions, which exhibit greater cytoarchitectural heterogeneity, thereby reducing the strength of structure–microstructure coupling observed in the TLE-SF cohort. Other features (*i.e.,* cell size, mean, variance) were not significant in either group (**Fig. S8**).

### Associations with gene expression

Network epicenters may preferentially emerge in regions with molecular profiles that facilitate seizure generation and propagation. To investigate this, we related patient-derived epicenter maps to regional gene expression patterns from the Allen Human Brain Atlas, fitting separate PLS models for TLE-SF and TLE-NSF ^34,35^(**Fig. 4a**). Cross-validation was used to select the optimal number of components, and spin permutation tests assessed statistical significance while accounting for spatial autocorrelation.

**Figure 4.**
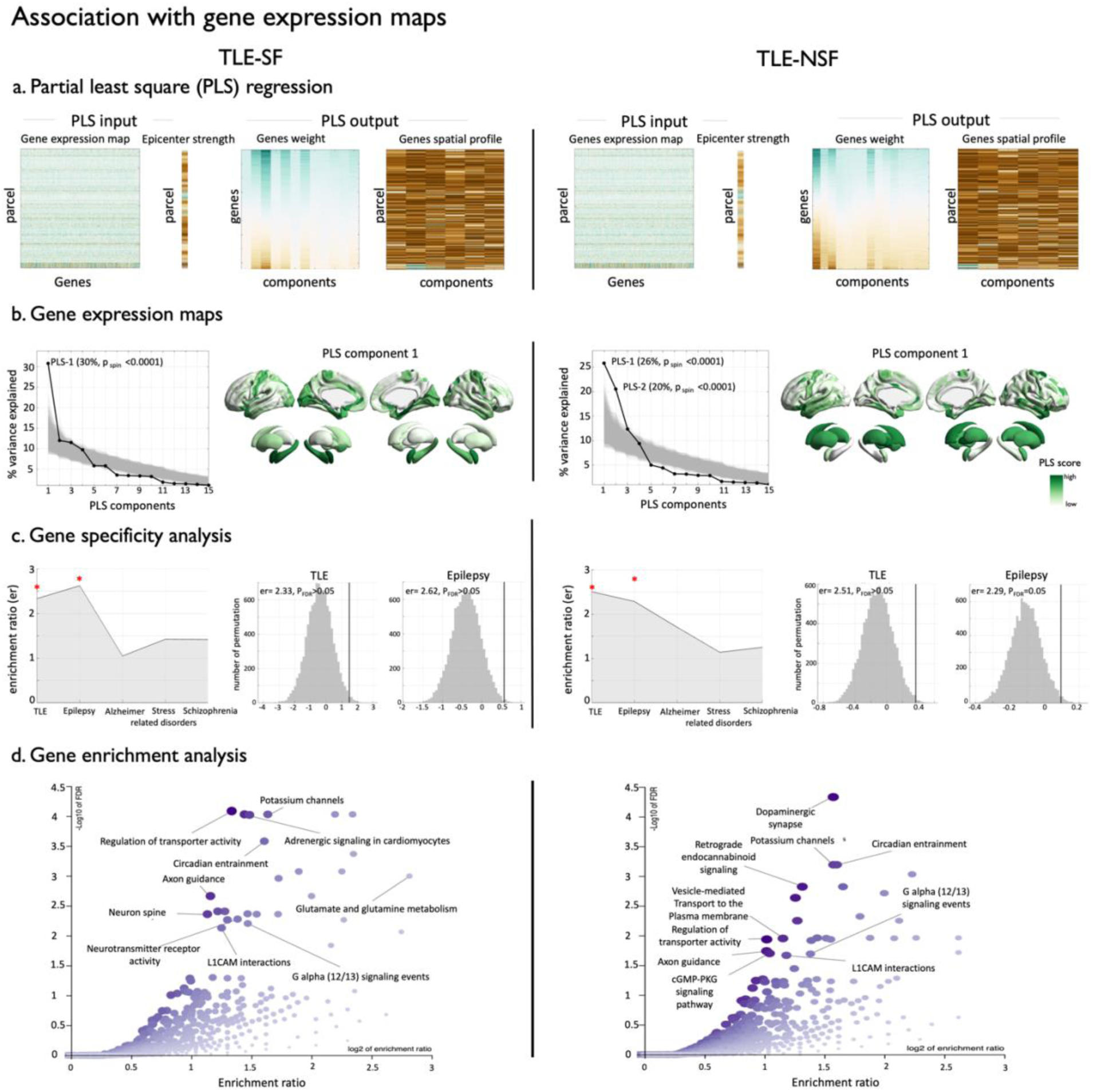
Association with gene expression maps. **(a)** Partial least squares (PLS) regression analysis in TLE-SF (left) and TLE-NSF (right) patients. Inputs were expression levels of 8,425 cortical and subcortical genes derived from the Allen Human Brain Atlas (AHBA) transcriptomic dataset. The response variable (Y) was defined as the mean structural and functional epicenter strength per region, averaged across cortical and subcortical measures and z-scored. **(b)** PLS identified significant components that maximized covariance between gene expression and epicenter strength. P_spin_ denotes significance after correcting for spatial autocorrelation. Brain maps showing gene expression patterns associated with significant components. **(c)** The plot shows the enrichment ratio of top-ranked genes for each surgical group across curated gene sets, including TLE, epilepsy, Alzheimer’s disease, stress-related disorders, and schizophrenia. The area of each bar reflects the magnitude of enrichment. Statistically significant enrichment (P_FDR_ < 0.05) is indicated by a red star above the corresponding bar. For significant gene, the corresponding null distribution from the permutation test is displayed to the right, with black lines indicating observed significant correlations. **(d)** Gene enrichment based on top 20% weighted genes (loadings).

One PLS component in TLE-SF and two components in TLE-NSF explained a substantial proportion of covariance between epicenter and gene expression maps. In TLE-SF, PLS-1 accounted for 30% of the covariance (P_spin_<0.0001). In TLE-NSF, PLS-1 and PLS-2 explained 26% and 20% of the covariance, respectively (P_spin_<0.000l, **Fig. 4b**). The spatial distribution of PLS score in TLE-SF was localized mostly to ipsilateral hippocampus, mesial temporal and fronto-central regions. In TLE-NSF one component was prominently localized to the hippocampus, whereas the other component was more widespread, encompassing fronto-central and lateral temporal cortices as well as subcortical regions, including the thalamus and basal ganglia. (**Fig. 4b**, **Fig. S9**).

We then asked whether the molecular signatures captured by the PLS components specifically reflect genetic risk for TLE and epilepsy. Using a gene specificity analysis, we compared the top-ranked PLS genes to curated lists of TLE- and epilepsy-associated genes, as well as genes linked to other hippocampus-related conditions (**Supplementary Table 1**). Both TLE-SF and TLE-NSF components were preferentially enriched for TLE- and epilepsy-related genes (P_FDR_ < 0.05), whereas genes associated with Alzheimer’s disease, stress-related disorders (depression, anxiety, PTSD), and schizophrenia showed no significant enrichment (**Fig. 4c**). These results indicate that the PLS-derived molecular patterns are more closely aligned with epilepsy-related mechanisms rather than reflecting general hippocampal pathology seen in other diseases.

Having identified the molecular components associated with network epicenters via PLS, we next asked which disease-related and biological pathways these components reflect and how they differ between TLE-SF and TLE-NSF. To address this, we performed gene set enrichment analysis on the top 20% of genes from PLS components, identifying over-represented pathways across multiple functional databases with FDR correction. Gene set enrichment analysis revealed distinct molecular profiles associated with surgical outcome in TLE. The TLE-SF group showed strong and more focused enrichment (size=41 genes; enrichment ratio=2.52; P_FDR_<0.0001) for calcium-dependent signaling and synaptic regulation. In contrast, TLE-NSF group showed broader enrichment (size=132 genes; enrichment ratio=2.98; P_FDR_<0.0001), reflecting widespread dysregulation of pathways involved in neuronal excitability, intracellular signaling, and neuromodulation. These results suggest that patients who achieve seizure freedom harbor a more coherent and functionally concentrated pathological substrate, whereas persistent seizures are associated with broader network-level molecular disturbances (**Fig. 4d**).

### Association with extent of resection

Finally, we investigated whether surgical targeting of network epicenters could help explain differences in postoperative outcomes. Using the top 10% of structural and functional epicenters, we quantified their overlap with resected tissue for each patient, accounting for age, sex, histopathology, total resection volume, and disease duration. **Fig. 5a** illustrates the probability of resection mapped in voxel space and on cortical surfaces for TLE-SF (*left*) and TLE-NSF (*right*) patients. The total volume of resection did not differ between TLE-SF and TLE-NSF patients (*t*=-0.52, *p*=0.60; **Fig. 5b, left**), indicating comparable surgical extents across groups. Despite this, TLE-SF patients showed significantly greater overlap of both structural (*t*=2.97, *p*=0.001; **Fig. 5b, middle**) and functional (*t*=11.52, *p*<0.001; **Fig. 5b, right**) epicenters with resected tissue compared to TLE-NSF cohort. These findings remained consistent after controlling for age, sex, histopathology, resection volume, and disease duration, and were robust across alternative top-parcel thresholds at the 20th, 30th, and 40th percentiles, highlighting that targeting key network epicenters rather than total resection extent may underlie favorable surgical outcomes.

**Figure 5.**
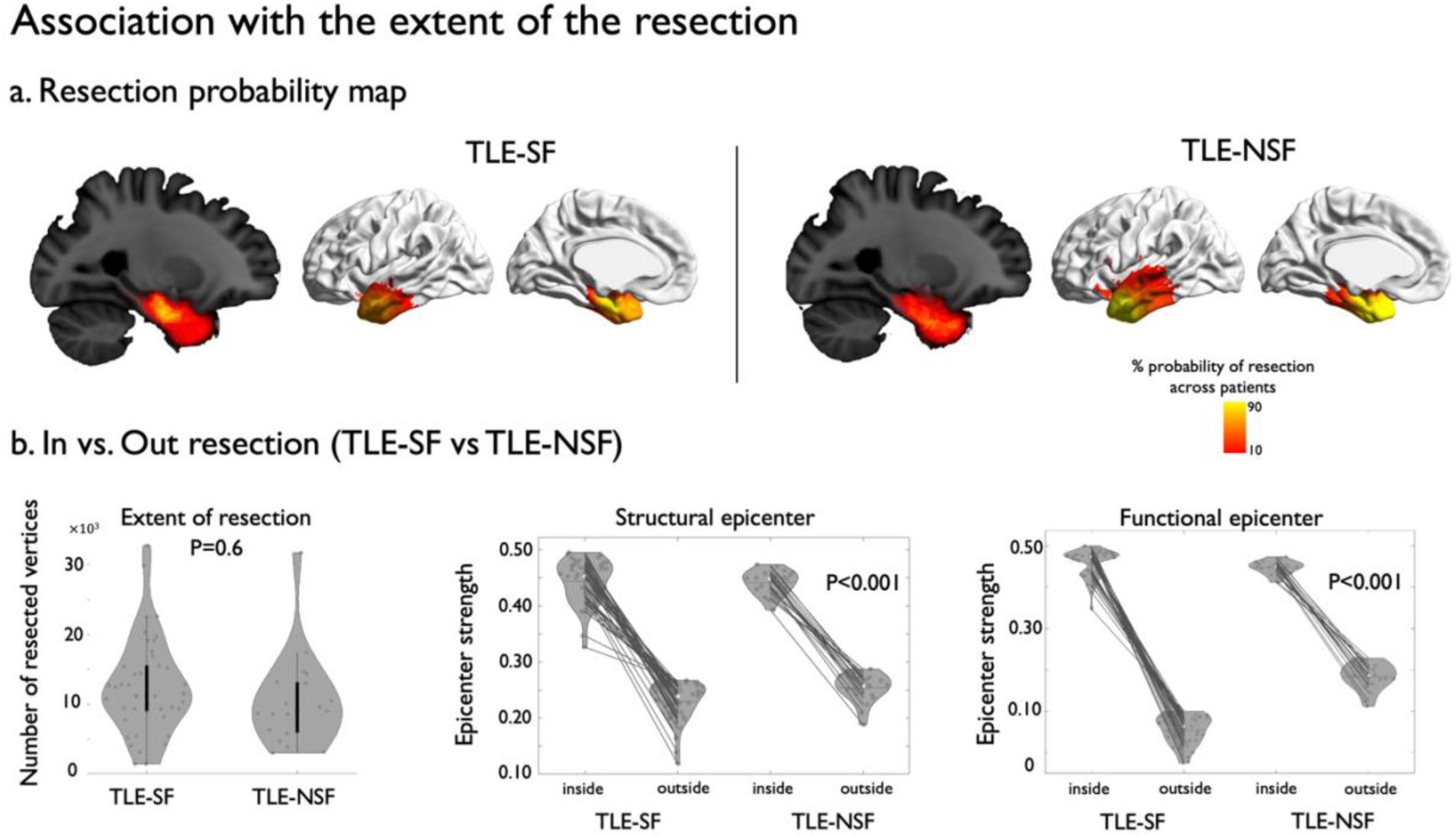
Association with the extent of resection. **(a)** Probability of resection masks mapped onto volumetric space (*left*) and cortical surface (*right*) for TLE-SF and TLE-NSF patients. **(b)** Violin plots comparing TLE-SF and TLE-NSF groups for (*left*) total resection volume, (*middle*) proportion of structural epicenters resected inside versus outside the cavity, and (*right*) proportion of functional epicenters resected inside versus outside the cavity.

## Discussion

This study introduces a multimodal precision neuroimaging framework to understand brain substrates that relate to post-surgical seizure outcomes in TLE. Moving beyond conventional group-level analyses, we quantify patient-specific deviations from healthy brain organization using a normative modelling framework. Moreover, we provide a multiscale contextualization of these findings, spanning connectome-based epicenter mapping, cytoarchitectural mapping, and transcriptomic profiling, to identify neurobiological substrates associated to surgical outcome in TLE. Our findings indicate that surgical failure likely reflects an interplay of molecular, cellular, and network level mechanisms that cannot be explained by the amount of resection alone. Residual epileptogenic activity within posterior mesiotemporal regions, extension into lateral temporal or insular cortices, and bilateral network engagement all contribute to seizure persistence. These divergent pathways are supported by distinct cytoarchitectural and molecular profiles, demonstrating that abnormalities across multiple biological scales converge to shape surgical success or failure. Collectively, our framework provides a unified lens on the multifactorial nature of epileptogenic networks and lays the foundation for individualized surgical planning, with molecular signatures emerging as potential prognostic biomarkers to guide more precise interventions.

Moving from traditional group-level analyses to an individualized normative modeling framework, we identified co-existing yet distinct axes of structural alterations in temporal lobe epilepsy that differentiate seizure-freedom from seizure recurrence after the surgery. Normative modeling provides a framework to quantify patient-specific deviations relative to healthy variation, capturing the unique patterns that define an individual’s pathology ^21^. By highlighting how abnormalities co-occur and contribute to clinically relevant outcomes, it offers a principled approach for linking biological deviations to patient-specific disease expression ^21–23^. Although TLE-SF and TLE-NSF groups exhibited a similar burden and topography of extreme deviations across structural and microstructural measures, our approach identified subtle, yet discernible regional nuances. TLE-SF patients exhibited spatially coherent abnormalities, characterized by anterior hippocampal atrophy accompanied by cortical thinning and fronto-temporal white matter disruption in adjacent association areas. These features are indicative of a focal mesiotemporal disease process largely contained within the resection boundaries. In contrast, TLE-NSF patients demonstrated more spatially scattered and bilateral deviations, encompassing posterior hippocampal regions and parahippocampal gyrus, ipsilateral sensory, and contralateral secondary somatosensory cortices, reflecting a broader and less hierarchically organized network pathology. Notably, in TLE-NSF, posterior hippocampal–parahippocampal abnormalities frequently extended beyond the conventional anterior temporal resection boundary, leaving abnormal tissue unresected. This likely reflects surgical restraint to minimize functional consequences ^36,37^. Interestingly, in a subset of patients with seizure recurrence, we observed a pattern of ipsilateral hippocampal and amygdala hypertrophy. Although rarely discussed, this phenomenon may reflect an active disease process rather than residual structural damage ^38^. Local gliotic or neuroinflammatory mechanisms could drive swelling, synaptic remodeling, and hyperexcitability, ^39^, sustaining an epileptogenic network post-resection ^7^. Alternatively, autoimmune or systemic inflammation may induce reactive hypertrophy via cytokine-mediated neuroplastic changes and increased excitatory signaling ^40–42^. Hippocampal–amygdala enlargement may thus serve as a biomarker of ongoing local or immune-mediated epileptic activity contributing to persistent seizures ^7,42,43^.

Findings were overall consistent in left and right TLE patients, with however subtle variations between groups. While left TLE-NSF patients showed pronounced alterations in the insular cortex, right TLE-NSF patients exhibited more marked deviations in associative regions, including the lateral temporal neocortex. In neither group did the hippocampus host the most marked deviations, providing evidence that surgical failure may stem from temporal-plus network involvement ^44,45^. This pattern suggests that in TLE-NSF, the hippocampus is structurally less affected and may instead act as a functionally engaged node within a broader epileptogenic network, rather than serving as the primary seizure generating region ^46,47^. This is also supported by our epicenter mapping findings where the hippocampus emerged as functionally but not structurally abnormal. Taken together, by quantifying individualized deviations from normative brain structure, this framework reveals hidden dimensions of disease heterogeneity in TLE. The distinct spatial signatures observed across outcome groups suggest that seizure freedom after surgery may relate to a more localized and structurally contained epileptogenic substrate, whereas post-operative seizure recurrence may reflect a more distributed and system-level pathology. This individualized perspective provides a biologically meaningful bridge between macroscopic imaging markers and the variable clinical trajectories seen in TLE.

Epicenter mapping provided further insight into the network organization underlying these morphometric patterns by identifying regions that act as the central nodes of distributed network pathology. This approach has proven valuable in clinical research by identifying network mechanisms that link regional abnormalities to system-level effects, and highlight regions that shape patient outcomes ^23,25,48^. From a clinical perspective, this shifts interpretation from isolated MRI findings to whether the surgically targeted region captures the core epileptogenic network, a distinction that is critical for explaining surgical success versus failure. In TLE-SF, structural and functional epicenters were spatially concordant and restricted to the ipsilateral mesial temporal system. This cohesive and structurally anchored configuration aligns with the regions typically removed in standard anterior temporal resections, supporting effective network disconnection and seizure freedom ^8,49^. In contrast, TLE-NSF patients showed weaker alignment between structural and functional epicenters. The hippocampus emerged as a functional but not structural epicenter, implying that it likely acts as an early relay node within a broader epileptogenic network rather than serving as the primary generator ^50,51^. Structural epicenters in these patients extended into lateral and posterior temporal cortices, which are regions often spared from resection. Functional epicenters additionally included contralateral homologues. This broader, distributed network likely underlies persistent seizures, reflecting contributions from residual mesiotemporal and extramesial tissue ^4,7,52^, contralateral recruitment ^53–55^, and temporal plus epilepsy ^44,56^ including insular involvement ^45,57^. Although MRI cannot directly trace ictal propagation, distributed epicenter patterns parallel stereo-encephalographic evidence suggesting that early extrahippocampal recruitment may relate to the risk of post-operative seizure recurrence ^46^. Together, these findings support the concept of a “temporal-plus” pathological network as one of the reasons underlying surgical failure in TLE.

Growing evidence across neurological and psychiatric disorders demonstrates that macroscale network abnormalities are not arbitrary, but are preferentially anchored to specific cytoarchitectural and transcriptional contexts, reflecting shared biological constraints on disease expression and propagation ^58,59^. Within this framework, contextualizing network epicenters using histological and molecular reference maps provides insights into how the intrinsic microstructural and molecular properties of these regions relate to clinical outcomes. Histological contextualization analyses revealed that structural-functional epicenters in TLE-SF patients preferentially localized to limbic and paralimbic cortices, regions characterized by thicker cortex, larger neurons, lower neuronal density, and increased laminar skewness. These microstructural features reflect evolutionarily older cortical types with relatively simple and coherent laminar architecture ^27,60^, suggesting that pathological activity in these patients is embedded within structurally and functionally constrained cortical territories. These cortices are arranged in a ring-like formation at the base of the cortex, display little laminar differentiation, and exhibit high plasticity ^26,27^. While increased plasticity can render them more vulnerable to pathology, such as mossy fiber sprouting and synaptic reorganization ^61,62^, their simpler laminar architecture may constrain epileptogenic activity within locally coherent networks, limiting widespread propagation and making focal resections more likely to disrupt the core epileptogenic zone ^63,64^. In contrast, epicenters in patients with persistent seizures were more widely distributed across neocortical regions, which exhibit greater cytoarchitectural heterogeneity. The more differentiated laminar structure and heterogeneous microcircuitry of these regions may facilitate broader network propagation, consistent with the distributed epicenters observed in TLE-NSF ^65,66^. Complementing microstructural observations, PLS analyses revealed that structural-functional epicenters corresponded to distinct molecular signatures. In TLE-SF, epicenters were preferentially enriched for calcium-dependent signaling pathways, suggesting focal dysregulation of neuronal membrane excitability and action potential dynamics. By contrast, TLE-NSF epicenters showed broader enrichment for pathways governing neuronal excitability, intracellular signaling, and neuromodulation, consistent with distributed alterations in network-level excitability. These molecular signatures converge with prior reports linking specific excitatory pathways to seizure persistence and post-surgical recurrence ^30,67^. Importantly, such gene expression profiles may have prognostic potential: as demonstrated in peripheral blood studies, specific mRNA and miRNA signatures can identify patient subgroups at higher risk for seizure recurrence after surgery ^67–69^. Similarly, our findings offer potential avenues for future work testing if the molecular composition of TLE epicenters could serve as a biomarker for persistent epileptogenic networks, able to inform predictive models for individualized surgical planning. Collectively, these results indicate that multiscale properties, notably macroscale network embedding, cytoarchitecture, and molecular milieu, shape the focality of epileptogenic tissue, thereby influencing surgical outcomes. Integrating preoperative imaging with surgical anatomy provides a framework to understand how the relationship between pathological regions and resection boundaries relates to postoperative outcomes. Previous studies have demonstrated that combining morphometric and network-based analyses with surgical maps can help explain variability in surgical success and provide insight into the mechanisms underlying seizure persistence ^7,10,70,71^. Building on these, our findings suggest that seizure freedom is associated with a surgery effectively captures the dominant network epicenters, which are embedded within a compact, ipsilateral mesiotemporal network with strong structural–functional coherence. On the other hand, persistent seizures occur when these core drivers are spared, despite resections being of similar overall volume. Importantly, this suggests that surgical failure is not simply due to insufficient resection but reflects incomplete capture of the dominant epileptogenic network core ^7,72,73^. From a clinical standpoint, our study offers a principled framework for treatment calibration by identifying patients in whom standard resection is sufficient, those who may benefit from extended or tailored resections, and those in whom neuromodulatory or network-targeted interventions may be more appropriate than further tissue removal. By shifting from a focus-centered to individualized network-centered model of epilepsy, this approach has the potential to refine electrode placement, optimize surgical boundaries, and guide personalized therapeutic strategies in drug-resistant TLE ^73,74^.

## Materials and methods

### Participants

We studied 102 pharmaco-resistant TLE patients (58 females, mean age ± SD = 32.6 ± 10.4 years, range=15-61) who underwent epilepsy surgery and had at least 1 year of postoperative follow-up as well as 94 age- and sex-matched healthy controls (45 female, 34.4 ± 10.4, range=19-65). Participants were selected from 3 independent datasets of 2 tertiary epilepsy centers: (i) Montreal Neurological Institute-Hospital (*MICA-MICs*: TLE/healthy controls = 11/23; *NOEL*: 51/40) (ii) Jinling Hospital (*Nanj*: 40/31). Patients were diagnosed according to the classification of the International League Against Epilepsy based on a comprehensive evaluation that included clinical history, seizure semiology, video-electroencephalography recordings, neuroimaging, and neuropsychological assessment. We excluded patients who had encephalitis, malformation of cortical development or a history of traumatic brain injury and those with a diagnosis of bilateral TLE. Patients were classified as seizure-free (TLE-SF, n = 77) or non-seizure-free (TLE-NSF, n = 25) based on post-surgical outcomes. The mean follow-up after the surgery was 4.9±2.5 years (range=1-10 years). Based on the histological analysis of the resected specimen ^75^, we observed a higher proportion of hippocampal cell loss and gliosis versus isolated gliosis in TLE-SF compared to TLE-NSF (n=66/16 *vs.* 11/9; χ^2^=4.35, p=0.03). No significant differences were found between groups with respect to the age at surgery (TLE-SF: 33.54 ± 10.6 years, range: 17–61; TLE-NSF: 29.56 ± 9.35 years, range: 15–45; *t* = 1.12, *p* = 0.2), sex distribution (TLE-SF: 42/77 female; TLE-NSF: 17/25 female; χ² = 0.6, *p* = 0.4), age at seizure onset (TLE-SF: 16.04 ± 11.06 years; TLE-NSF: 16.52 ± 9.19 years; *t* = 0.003, *p* = 0.9), or seizure lateralization (TLE-SF: 37/77 left; TLE-NSF: 11/25 left; χ² = 0.3, *p* = 0.5). TLE-SF patients showed a non-significant trend toward longer disease duration compared to TLE-NSF patients (TLE-SF: 17.49 ± 11.82 years; TLE-NSF: 13.04 ± 8.55 years; *t* = 1.58, *p* = 0.1). The study was approved by the research ethics boards of all participating sites, and all participants provided written informed consent in accordance with the Declaration of Helsinki.

### MRI acquisition and preprocessing

Multimodal MRI data, including T1-weighted, diffusion MRI, and rs-fMRI, were acquired with 3T scanners (*MICA-MICs*: Siemens Prisma; *Nanj* and *NOEL*: Siemens Trio) in all individuals (prior to surgery for patients). Multimodal MRI preprocessing was performed using micapipe, an open-access image processing and data fusion pipeline [v0.2.0; https://micapipe.readthedocs.io; ^76^]. Full details on MRI data acquisition and processing are provided in the Supplementary Materials. In brief, native T1w structural images were deobliqued, reoriented to standard neuroscience orientation, corrected for intensity nonuniformity, intensity normalized, skull-stripped and submitted to FreeSurfer [v6.0; https://surfer.nmr.mgh.harvard.edu; ^77^] to extract models of the cortical surfaces. Subcortical structures were segmented using FSL FIRST ^78^. Native diffusion weighted images (DWI) were denoised, underwent b0 intensity normalization and were corrected for susceptibility distortion, head motion and eddy currents. The hippocampus was segmented using a surface-based approach which allows inter-individual alignment of topologically homologous tissues in a flat-mapped 2D space, using HippUnfold [v0.2.3 https://hippunfold.readthedocs.io/; ^79,80^].

### Surgical cavity delineation

For 65 cases with available postoperative MRI (mean interval from surgery to post-op scan: 2.43 ± 3.15 years), resection cavities were manually segmented on each subject’s postoperative T1-weighted scan using ITK-SNAP. The resulting binary cavity masks were rigidly registered to the corresponding preoperative T1-weighted image to ensure accurate alignment in native space. Cavity masks were then projected onto the fsLR-32k pial, mid-thickness, and white cortical surfaces, and subsequently combined into a union mask to capture the full resection extent across cortical depths. All segmentations, registrations, and surface mappings were visually inspected to confirm accuracy.

## Statistical analysis

### Normative modeling analysis (Individualized *w*-score maps)

To account for potential variations across scanners and datasets, we applied ComBat harmonization to each MRI features independently ^81^. ComBat uses multivariate linear mixed-effects regression with empirical Bayes estimation to correct for batch effects while preserving biologically meaningful variability. The harmonized data were then used for all subsequent analyses.

We examined complementary structural imaging features indexing cortical morphology and white matter microstructure, including CT, FA, and ADC. Together, these measures capture distinct but related aspects of gray matter integrity and white matter organization in TLE. For each feature, surface-based values were averaged within cortical regions defined by the Glasser 360 parcellation ^82^. W-scores were calculated for each cortical parcel and for seven mesiotemporal and subcortical structures (*i.e.,* hippocampus, amygdala, caudate nucleus, putamen, thalamus, nucleus accumbens, globus pallidus) relative to the healthy control reference group ^83^. Analogous to a z-score (mean = 0, SD = 1), the w-score quantifies the deviation of an individual’s observed value from the normative expectation, adjusted for covariates such as age and sex. Specifically, w-scores were computed as *w* score = (observed – expected) / residual standard deviation where *Observed* is the participant’s MRI metric, *Expected* is the predicted value from the normative model, and *Residual SD* is the standard deviation of model residuals. Thus, w-scores represent z-scores corrected for demographic covariates ^22,23,83^. Right TLE cases were flipped along the midline to yield ipsilateral and contralateral representations.

Deviations from normative distributions were estimated for each feature. Parcels with |W-scores| ≥ 1.96 were classified as extreme deviations. For each patient group (TLE-SF *vs*. TLE-NSF), we generated abnormality probability maps by computing the proportion of patients with significant decreases (CT, FA) or increases (ADC) at each parcel, across both cortical and subcortical regions. Results were subsequently stratified by cytoarchitectonic classes defined in the Von Economo and Koskinas parcellation ^84^. To integrate information across modalities, we computed a composite multivariate score for each individual by estimating the Mahalanobis distance of the joint distribution of CT, FA, and ADC. Extreme deviations of the multivariate w-score were similarly thresholded (|w| ≥ 1.96) and visualized to summarize patient-specific structural abnormalities. Finally, differences between TLE-SF and TLE-NSF were assessed using linear models. The multivariate score provided a summary measure of structural deviations for subsequent analysis. In addition, we: (i) performed surface-based assessments of hippocampal w-scores across all features (CT, FA, ADC) as well as the multivariate w-score, (ii) compared TLE-SF *vs* TLE-NSF patients within right and left TLE separately, (iii) performed subgroup comparisons within HS and gliosis groups, and (iv) evaluated cross-site consistency by repeating analyses within individual acquisition sites.

### Disease epicenter mapping

To identify candidate network epicenters associated with widespread abnormalities in TLE-SF and TLE-NSF, we employed a seed-based correlation framework. Normative structural and functional connectivity matrices were constructed from healthy control participants in our dataset by averaging streamline-weighted structural connectivity and resting-state functional connectivity, respectively. All connectivity matrices were parcellated using the Glasser-360 atlas with additional subcortical regions. For each cortical and subcortical parcel, we extracted its whole-brain connectivity profile and correlated this profile with the spatial pattern of patient-specific multivariate abnormality maps (i.e., multivariate w-score maps). This procedure yielded an “epicenter strength” value, quantifying how well a region’s connectivity pattern aligned with the observed abnormality distribution ^23–25^. Spatial significance of cortical epicenters was assessed using spin permutation tests (1,000 rotations) to control for spatial autocorrelation, whereas mesiotemporal and subcortical epicenters were evaluated using standard permutation-based correlation tests. Analyses were conducted separately for structural and functional networks and for TLE-SF and TLE-NSF patient subgroups. As an additional step, we compared TLE-SF and TLE-NSF epicenter strength by computing the observed difference for each parcel (TLE-SF minus TLE-NSF) and testing significance against the same null distributions. Parcels exceeding a threshold of P_FDR_ < 0.05 were retained to generate a group-difference map highlighting regions with relatively stronger epicenter involvement in TLE-SF versus TLE-NSF. Significant epicenters were visualized on cortical and subcortical surface templates. In addition, we: (i) repeated the epicenter identification analyses while using independent healthy control dataset (ii) evaluated cross-site consistency by repeating analyses within individual acquisition sites.

### Association with cytoarchitecture maps

To examine associations between patient-derived epicenter maps and cytoarchitectural organization, we used two independent cytoarchitectural reference maps. The first was the von Economo–Koskinas MRI atlas, which provides quantitative measures of cell size, cell density, and cortical thickness across 43 cortical regions per hemisphere [http://dutchconnectomelab.nl; ^85^].The second was the BigBrain atlas, a high-resolution 3D reconstruction of a stained postmortem human brain, downloaded via the ENIGMA Toolbox [https://enigma-toolbox.readthedocs.io; ^86,87^]. From the BigBrain dataset, we extracted the first three statistical moments of intracortical intensity profiles—mean (overall cell density), variance (laminar differentiation), and skewness (balance between upper and deeper layers). Cytoarchitectural features from both atlases were mapped onto intracortical surface models in standard space and aligned to the Glasser 360 parcellation.

Regional epicenter strength was defined as the mean of structural and functional epicenter centrality per parcel, averaged across cortical and subcortical regions separately for the TLE-SF and TLE-NSF groups. Associations between cytoarchitectural features and regional epicenter strength were assessed using spin permutation tests (1,000 rotations) to control for spatial autocorrelation.

### Associations with gene expression maps

We investigated the association between patient-derived epicenter maps and regional gene expression patterns using partial least squares analysis (PLS). Regional gene expression maps were downloaded from the ENIGMA Toolbox ^87^ which were sourced from the Allen Human Brain Atlas (AHBA)^88^ via *abagen* ^34^ in accordance with established recommendation ^35^. Briefly, probes were filtered and collapsed to genes, normalized within donors, and aggregated across parcels and donors. Genes with low cross-donor consistency (r<0.2) were excluded, which resulted in 8,425 genes being mapped to Glasser-360 cortical parcels and subcortical structures. All maps were z-scored prior to analysis. The response variable (*Y*) was defined as regional epicenter strength, computed as the mean of structural and functional epicenter centrality per parcel, averaged across cortical and subcortical regions and z-scored. PLS regression was then performed to identify molecular patterns associated with epicenter strength. Models were fit with a maximum of 20 components, with 10-fold cross-validation used to select the optimal number of components based on minimum cross-validated mean squared error. Component signs were aligned to maximize positive correlation with patient maps, and the variance explained in *X* and *Y* was computed. Statistical significance was assessed using 10,000 spin permutations to account for spatial autocorrelation, whereby cortical maps were permuted on fsaverage5 spheres and subcortical parcels shuffled independently. To evaluate stability, 10,000 bootstrap resamples were performed to estimate z-scores for each gene’s contribution. For significant components, genes were ranked by absolute loadings, and the top contributors were reported. Separate PLS models were fit for TLE-SF and TLE-NSF epicenter maps.

### Enrichment analysis

We performed gene set enrichment analysis using the WebGestalt toolkit (https://webgestalt.org) ^89^. For each latent variable, genes were ranked by their PLS-derived bootstrap weights, and the top 20% (based on absolute loadings) were selected. Over-representation analysis (ORA) was then performed using multiple functional databases, including Gene Ontology (biological process, cellular component, and molecular function), Reactome, and KEGG pathways. Enrichment was quantified as the ratio of observed to expected gene overlap, with statistical significance determined using hypergeometric testing and false discovery rate (FDR) correction implemented in WebGestalt.

### Gene specificity analysis

We tested whether genes associated with TLE ^90–92^, and epilepsy ^93,94^ were preferentially represented in the TLE-SF and TLE-NSF PLS components, compared with genes linked to other hippocampus-related conditions, including Alzheimer’s disease ^95,96^, stress-related disorders (depression, anxiety, PTSD) ^97,98^, and schizophrenia ^99^. The list of gene candidates is presented in **Supplementary table 1**. Enrichment ratios were calculated as the difference between the mean bootstrap weight of candidate genes and that of an equal number of randomly permuted genes, divided by the standard deviation of the permuted weights. Significance was assessed using 10,000 permutations. This analysis evaluates whether the spatial patterns captured by the PLS components specifically reflect genetic risk for TLE and epilepsy.

### Association with extent of resection

To investigate the association between resected tissue and regions of strongest abnormalities, we first performed a direct group comparison of total resection volume between TLE-SF and TLE-NSF patients. Resection volume was defined as the sum of vertices classified as resected within each patient’s surgical cavity, providing a vertex-wise estimate of surgical extent. We then assessed the degree to which regions of strongest network abnormality were selectively included within the resection. Specifically, we identified the top 10% of structural and functional epicenters using group-specific percentile thresholds. For each patient, epicenter strength was averaged separately for vertices within the resection cavity (“in-cavity”) and for vertices outside the cavity but within the top-parcel mask (“out-of-cavity”). Linear models were used to compare these measures between TLE-SF and TLE-NSF patients, controlling for age, sex, total resection volume, histopathological diagnosis, and disease duration. To ensure robustness, analyses were repeated using alternative epicenter thresholds (20th, 30th, and 40th percentiles).

## Data availability

The *MICA-MICs* dataset is available at the Open Science Framework (https://osf.io/j532r/) and the Canadian Open Neuroscience Platform (https://portal.conp.ca/). The *Nanj* and *NOEL* datasets will be made available upon reasonable request.

## Supplementary Method

### MRI Acquisition

*MICA-MICs dataset*. Data were collected on a 3-T Siemens Magnetom Prisma-Fit scanner equipped with a 64-channel head coil, and included: (i) two T1-weighted scans (3D-MPRAGE, repetition time [TR] = 2300 ms, echo time [TE] = 3.14 ms, flip angle [FA] = 9°, field of view [FOV] = 256×256 mm2, voxel size = 0.8×0.8×0.8 mm3, matrix size = 320×320, 224 slices), (ii) a resting-state functional MRI (fMRI) scan (multiband accelerated 2D-BOLD echo-planar imaging (EPI), TR = 600 ms, TE = 30 ms, FA = 52°, FOV = 240×240 mm2, voxel size = 3×3×3 mm3, multi-band factor = 6, 48 slices, 700 volumes), and (iii) a multi-shell diffusion MRI scan (2D spin-echo EPI, TR = 3500 ms, TE = 64.40 ms, FA = 90°, FOV = 224×224 mm2, voxel size = 1.6×1.6×1.6 mm3, 3 b0 images, b-values = 300/700/2000 s/mm2 with 10/40/90 diffusion directions). During the resting-state fMRI acquisition, participants were instructed to stay still, fixate a cross presented on the screen, and not to fall asleep.

*Nanj dataset*. Data were collected on a 3-T Siemens Trio scanner equipped with a 32-channel head coil, and included: (i) a T1-weighted scan (3D-MPRAGE, TR = 2300 ms, TE = 2.98 ms, FA = 9°, FOV = 256×256 mm2, voxel size = 0.5×0.5×1 mm3), (ii) a resting-state fMRI scan (2D gradient-echo EPI, TR = 2000 ms, TE = 30 ms, FA = 90°, FOV = 240×240 mm2, voxel size = 3.75×3.75×4 mm3, 30 slices, 255 volumes), and (iii) a diffusion MRI scan (2D spin-echo EPI, TR = 6100 ms, TE = 93 ms, FA = 90°, FOV = 240×240 mm2, voxel size = 0.94×0.94×3 mm3, 4 b0 images, b-value = 1000 s/mm2, 120 diffusion directions). During the resting-state fMRI acquisition, participants were instructed to keep their eyes closed and not to fall asleep.

*NOEL dataset*. Data were collected on a 3-T Siemens Trio scanner equipped with a 32-channel head coil, and included: (i) a T1-weighted scan (3D-MPRAGE, TR = 2300 ms, TE = 2.98 ms, FA = 9°, voxel size = 1×1×1 mm3), (ii) a resting-state fMRI scan (2D gradient-echo EPI, TR = 2020 ms, TE = 30 ms, FA = 90°, voxel size = 4×4×4 mm3, 34 slices, 150 volumes), and (iii) a diffusion MRI scan (2D twice-refocused EPI, TR = 8400 ms, TE = 90 ms, FA = 90°, voxel size = 2×2×3 mm3, 63 slices, 1 b0 images, b-value = 1000 s/mm2, 64 diffusion directions). During the resting-state fMRI acquisition, participants were instructed to keep their eyes closed and not to fall asleep.

### MRI preprocessing

MRI processing was carried out using micapipe (http://micapipe.readthedocs.io/), an open-access multimodal processing pipeline that integrates workflows for structural MRI, rs-fMRI, diffusion MRI, and myelin-sensitive MRI, enabling cross-modal feature integration^1^. In brief, *micapipe* automatically segmented subcortical and cortical regions using FreeSurfer^2^ (https://surfer.nmr.mgh.harvard.edu/fswiki) and FSL-FIRST^3^ on T1w MRI. Cortical thickness was computed as the Euclidean distance between corresponding points on the inner and outer cortical surfaces and mapped to the Glasser 360 atlas ^4^. Diffusion images were processed using MRtrix^5^ (https://www.mrtrix.org/), incorporating denoising, and correction for susceptibility distortions, head motion and eddy currents. As in prior studies ^6,7^, we generated a superficial white matter (SWM) surface running approximately 2mm below the grey-white matter boundary. This depth was chosen to target both the U-fiber system and terminations of long-range bundles that lie approximately between 1.5 and 2.5mm below the cortical interface. Fractional anisotropy (FA) and mean diffusivity (MD), proxies for fiber architecture and microstructure, were interpolated onto this SWM surface and then mapped to the Glasser 360 atlas. rs-fMRI data were processed using FSL ^3^ (https://fsl.fmrib.ox.ac.uk/fsl/fslwiki/) and AFNI^8^ (https://afni.nimh.nih.gov/), including discarding the first five volumes, reorientation, motion and distortion correction, skull stripping, removal of nuisance signals (*i.e*., white matter, cerebrospinal fluid) and artefacts using an ICA-FIX classifier^9^, and regression of high-motion timepoints. Volumetric timeseries were registered to native FreeSurfer space using boundary-based registration^10^ and mapped to participant-specific surface models using trilinear interpolation. Resulting surface-based timeseries were resampled to fslLR32, and downsampled to 5k vertices per hemisphere for computational efficiency. Participant-specific subcortical and hippocampal timeseries were non-linearly registered to corresponding native rs-fMRI space using ANTs^11^ (http://stnava.github.io/ANTs/), and averaged within the corresponding co-registered segmentations. The hippocampus was segmented using a surface-based approach which allows inter-individual alignment of topologically homologous tissues in a flat-mapped 2D space, using *HippUnfold* (https://hippunfold.readthedocs.io/)^12^. This tool incorporates a state-of-the-art U-Net deep learning framework and topologically-constrained subfield labelling^13,14^, and has shown excellent accuracy in hippocampal unfolding and subfield segmentation in healthy populations^13,15^.

---

**Figure S1.**
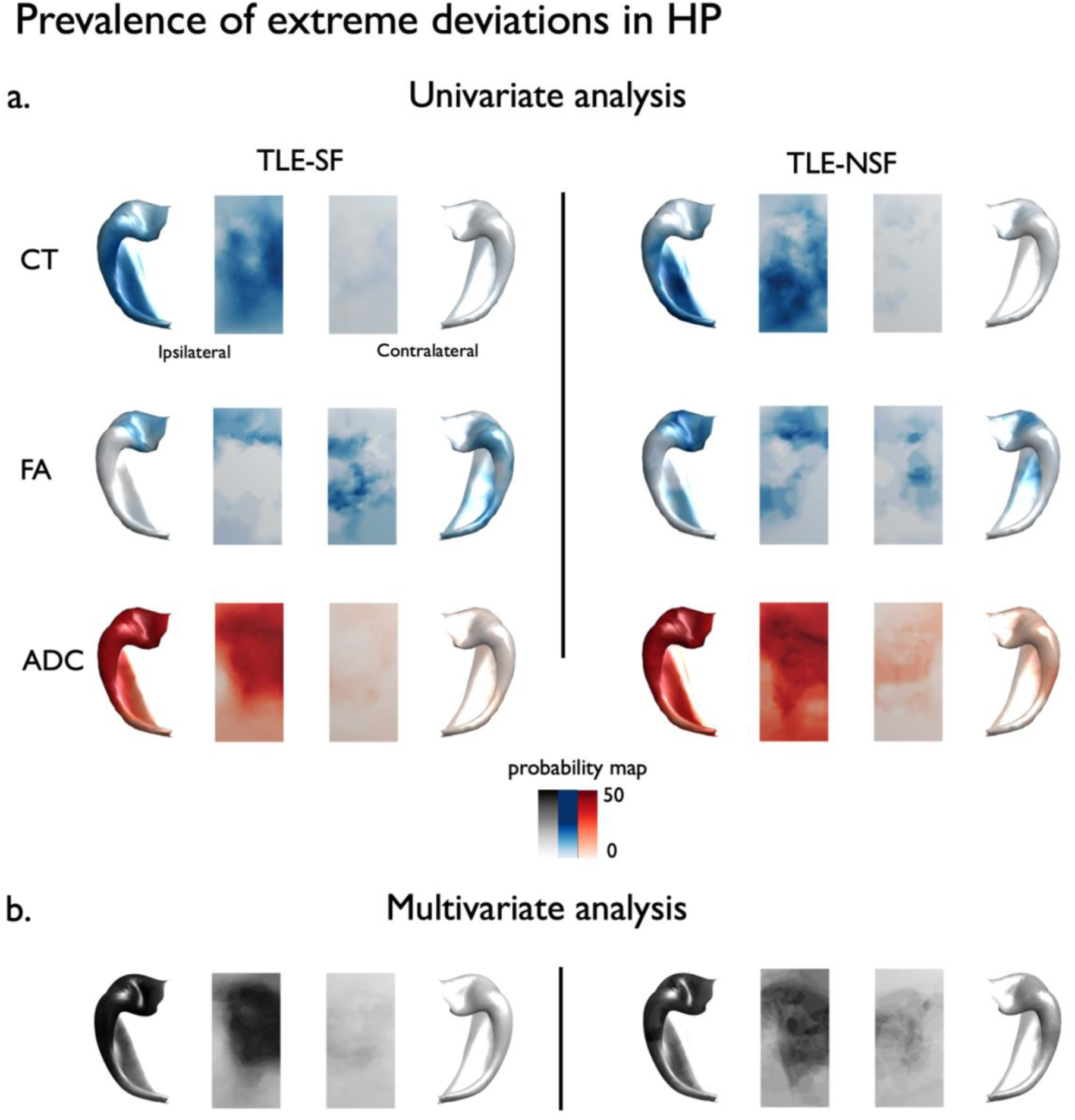
Prevalence of extreme deviations in hippocampus. **(A)** Proportion of TLE patients showing extreme deviations (|W-score| ≥ 1.96) in cortical thickness (CT), fractional anisotropy (FA), and apparent diffusion coefficient (ADC), displayed separately for seizure-free (TLE-SF, left) and non–seizure-free (TLE-NSF, right) groups. **(B)** Proportion of TLE-SF (left) and TLE-NSF (right) patients with multivariate W-scores (combined CT, FA, and ADC) exceeding ±1.96.

**Figure S2.**
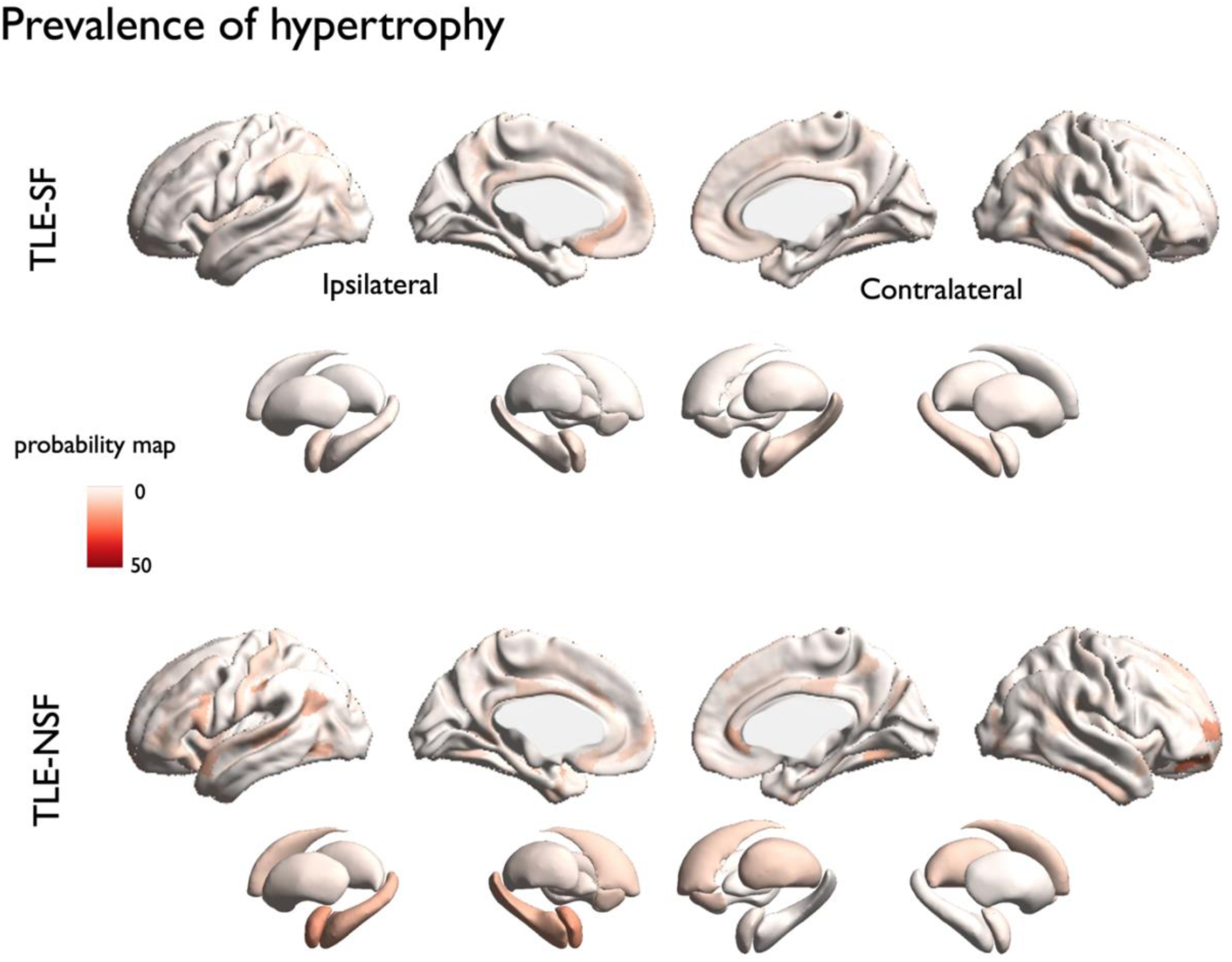
Prevalence of hypertrophy. Proportion of TLE patients showing hypertrophy (W-score ≥ 1.96) in cortical thickness (CT) displayed separately for seizure-free (TLE-SF, upper panel) and non–seizure-free (TLE-NSF, bottom panel) groups.

**Figure S3.**
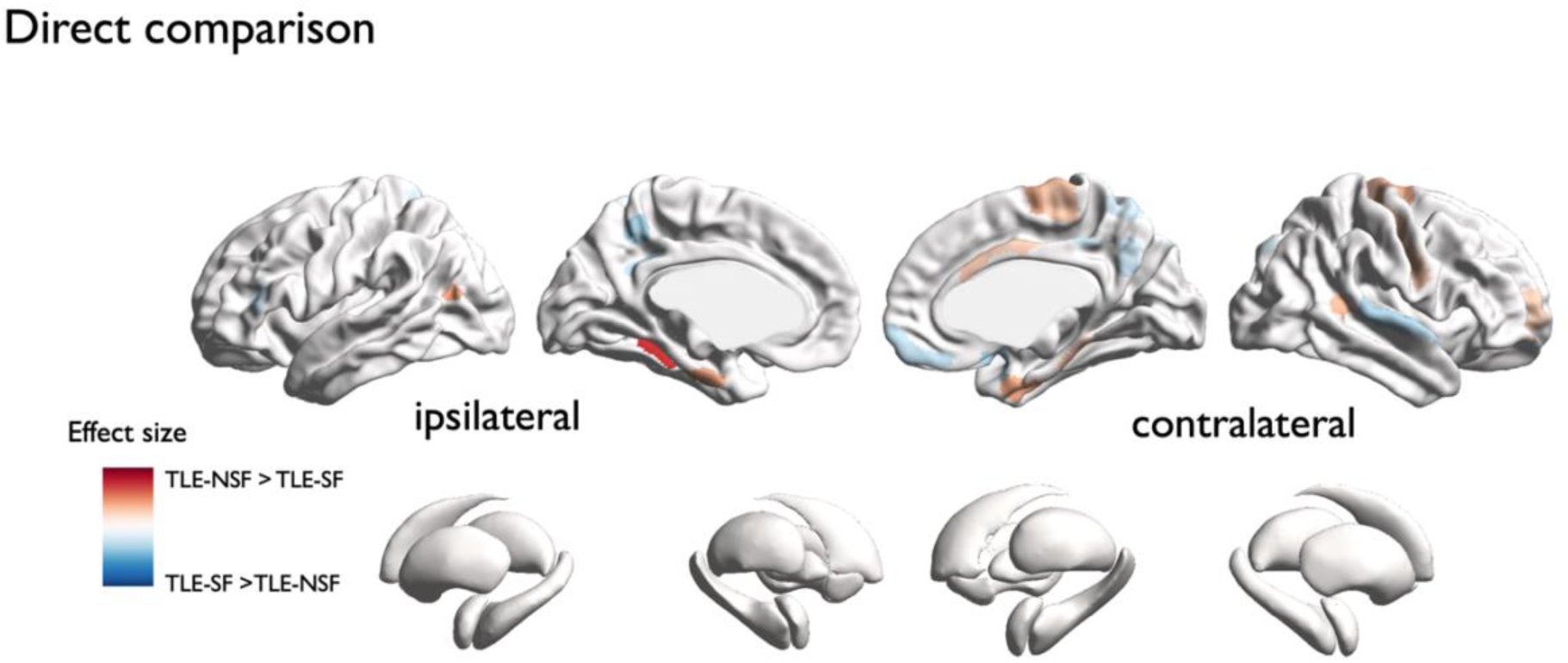
Direct comparison of multivariate w-scores between TLE-SF and TLE-NSF. Red indicates regions where TLE-NSF patients exhibit greater abnormalities, while blue indicates regions where TLE-SF patients show greater abnormalities.

**Figure S4.**
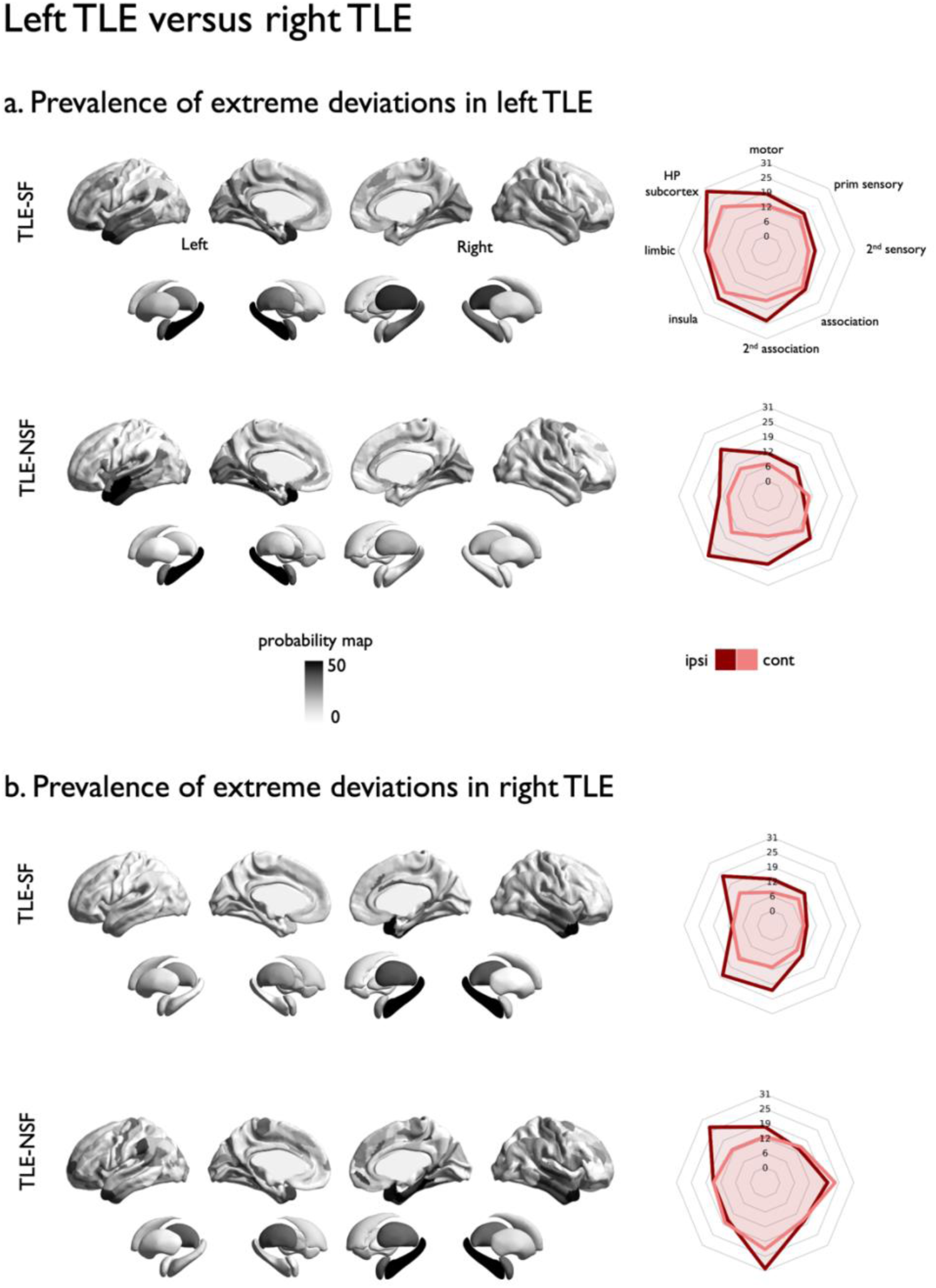
Prevalence of extreme deviations in left versus right TLE. Proportion of patients with seizure freedom (TLE-SF; top) and without seizure freedom (TLE-NSF; bottom) showing multivariate W-scores (combining cortical thickness, FA, and ADC) exceeding ±1.96. Results are displayed separately for left TLE (A) and right TLE (B). Spider plots illustrate the mean proportion of extreme multivariate deviations stratified by Von Economo cortical classes.

**Figure S5.**
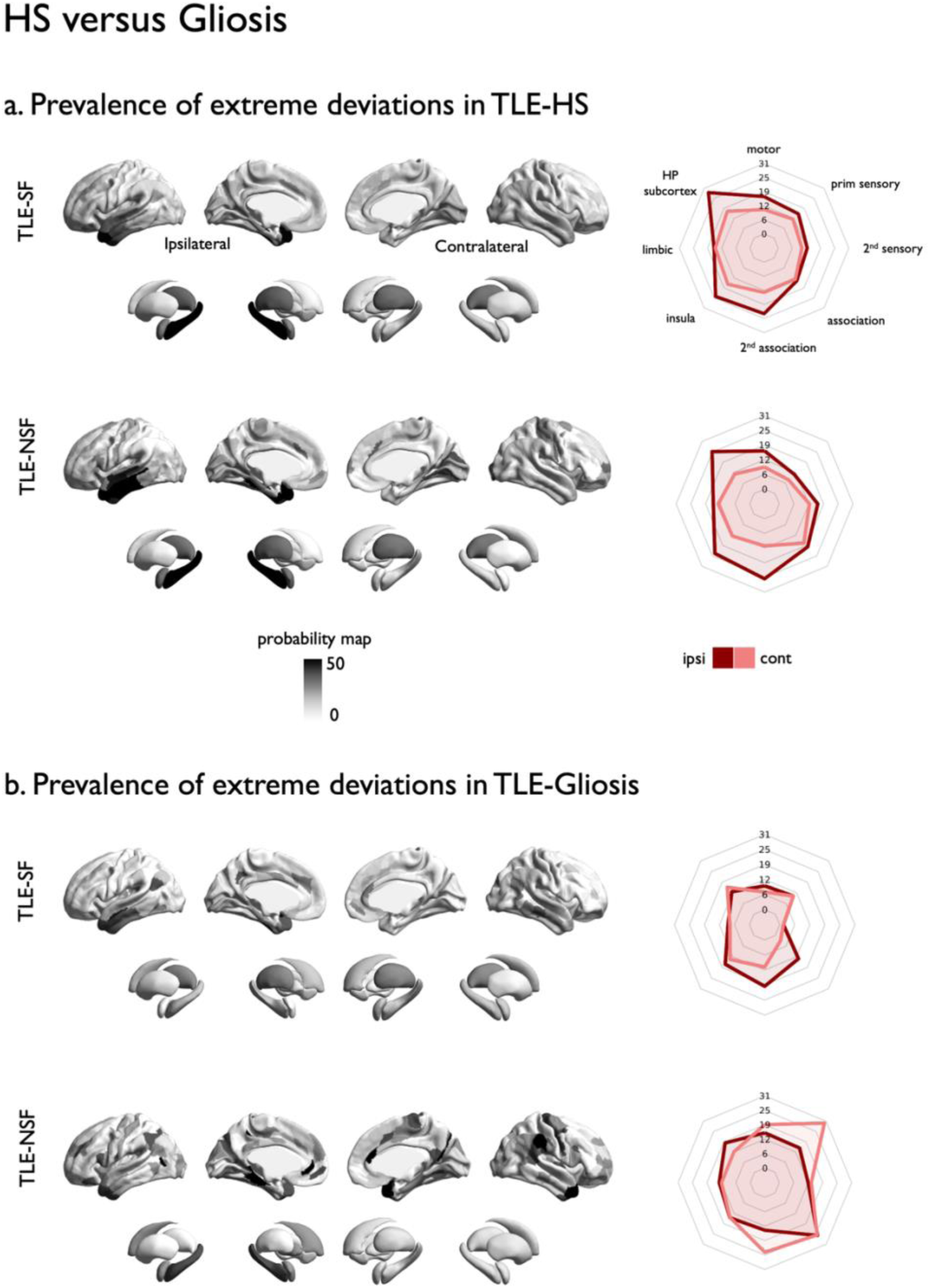
Prevalence of extreme deviations in HS versus Gliosis. Proportion of patients with seizure freedom (TLE-SF; top) and without seizure freedom (TLE-NSF; bottom) showing multivariate W-scores (combining cortical thickness, FA, and ADC) exceeding ±1.96. Results are displayed separately for TLE-HS (A) and TLE-gliosis (B). Spider plots illustrate the mean proportion of extreme multivariate deviations stratified by Von Economo cortical classes.

**Figure S6.**
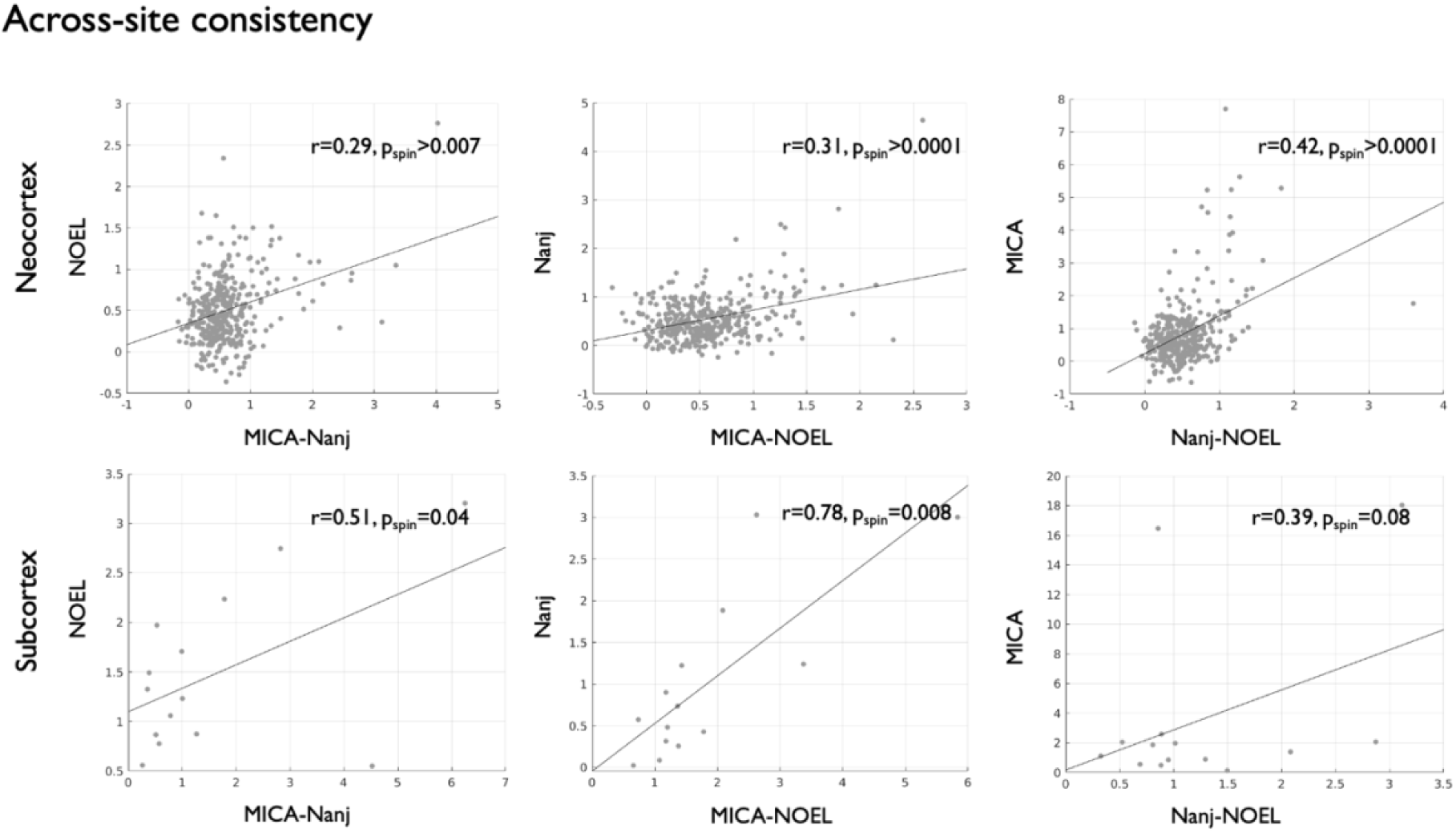
Across site consistency. Top: neocortex; Bottom: subcortex. For each site, scatterplots show region-wise multivariate w-scores from that site (y-axis) against the pooled mean of all *other* sites (x-axis). Pearson correlation significance was assessed with a spin test (10,000 rotations) to account for spatial autocorrelation; cortical maps used surface-based spherical rotations, and subcortical maps used within-hemisphere label permutations. Solid line indicates the least-squares fit.

**Figure S7.**
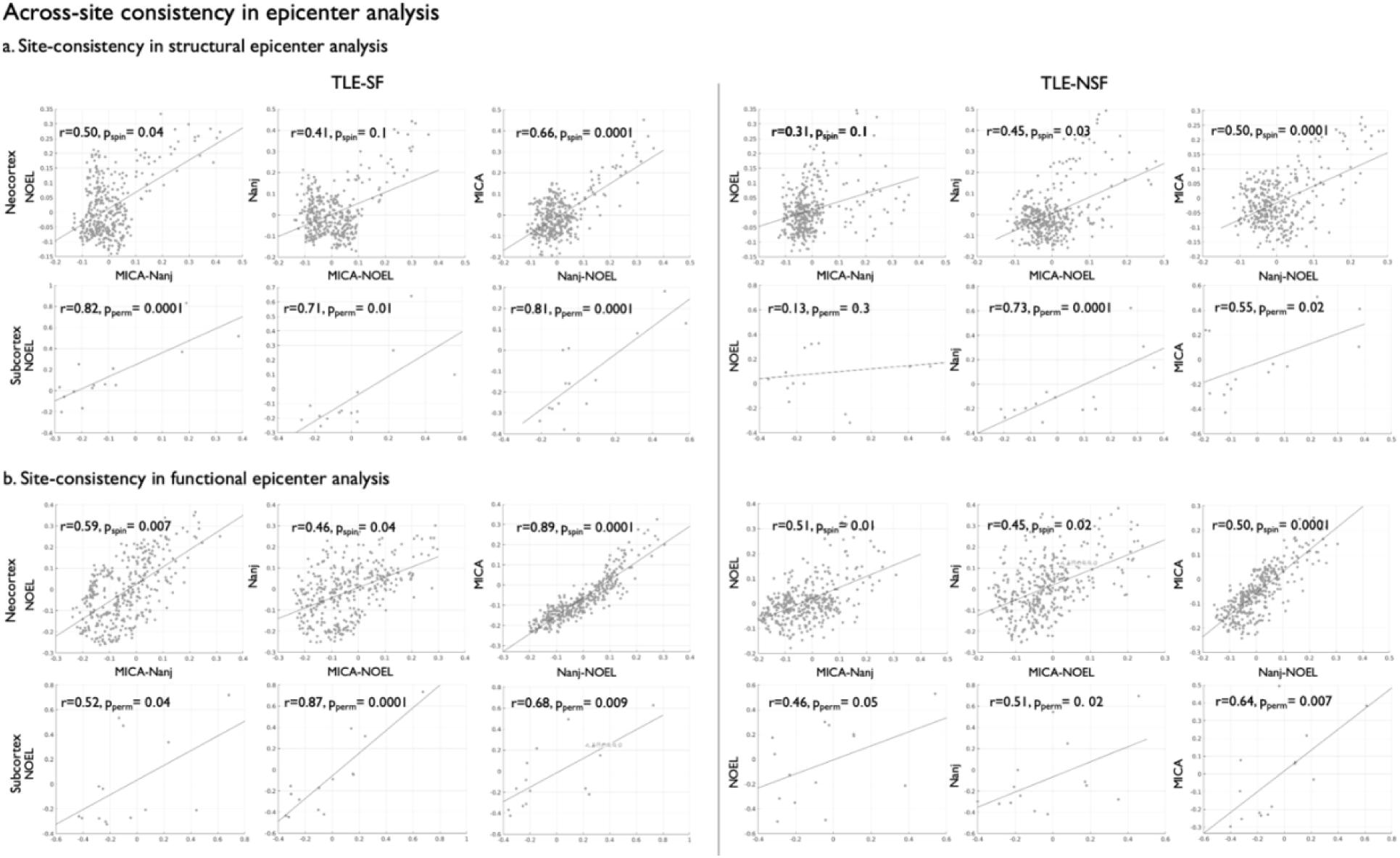
Across-site consistency in epicenter analysis. **a & b)** Top: neocortex; Bottom: subcortex. For each site, scatterplots show region-wise multivariate w-scores from that site (y-axis) against the pooled mean of all *other* sites (x-axis). Pearson correlation significance was assessed with a spin test (10,000 rotations) to account for spatial autocorrelation; cortical maps used surface-based spherical rotations, and subcortical maps used within-hemisphere label permutations. Solid line indicates the least-squares fit.

**Figure S8.**
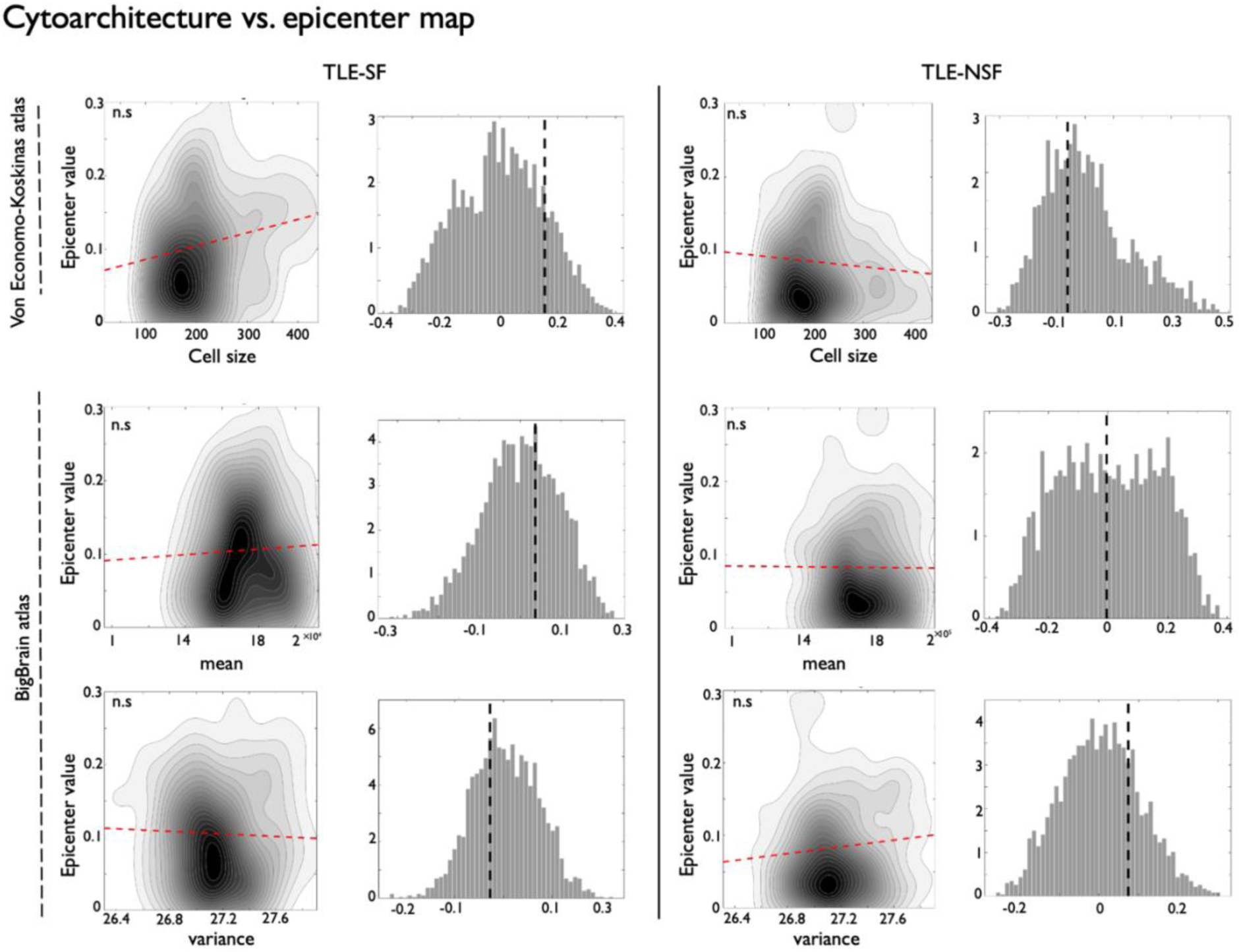
Association with cytoarchitecture maps. Contour plots show correlations between cytoarchitectural features and epicenters for TLE-SF (left) and TLE-NSF (right). For each contour plot, the corresponding null distribution from the spin test is displayed to the right, with dashed black lines indicating observed significant correlations.

**Figure S9.**
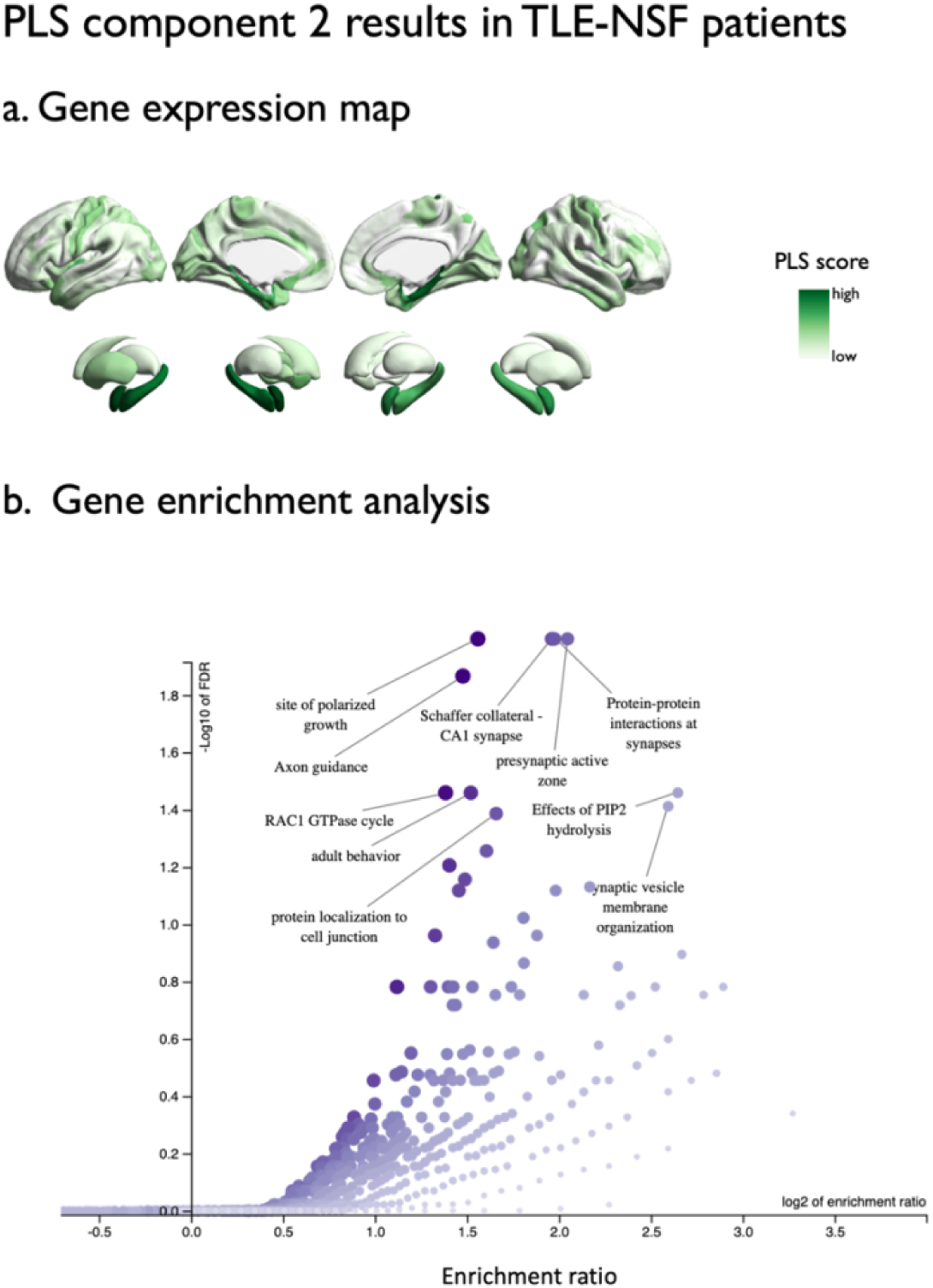
PLS component 2 results in TLE-NSF patients. (a) Brain maps showing gene expression patterns associated with significant components. (b) Gene enrichment based on top 20% weighted genes (loadings).

**Table S1.**
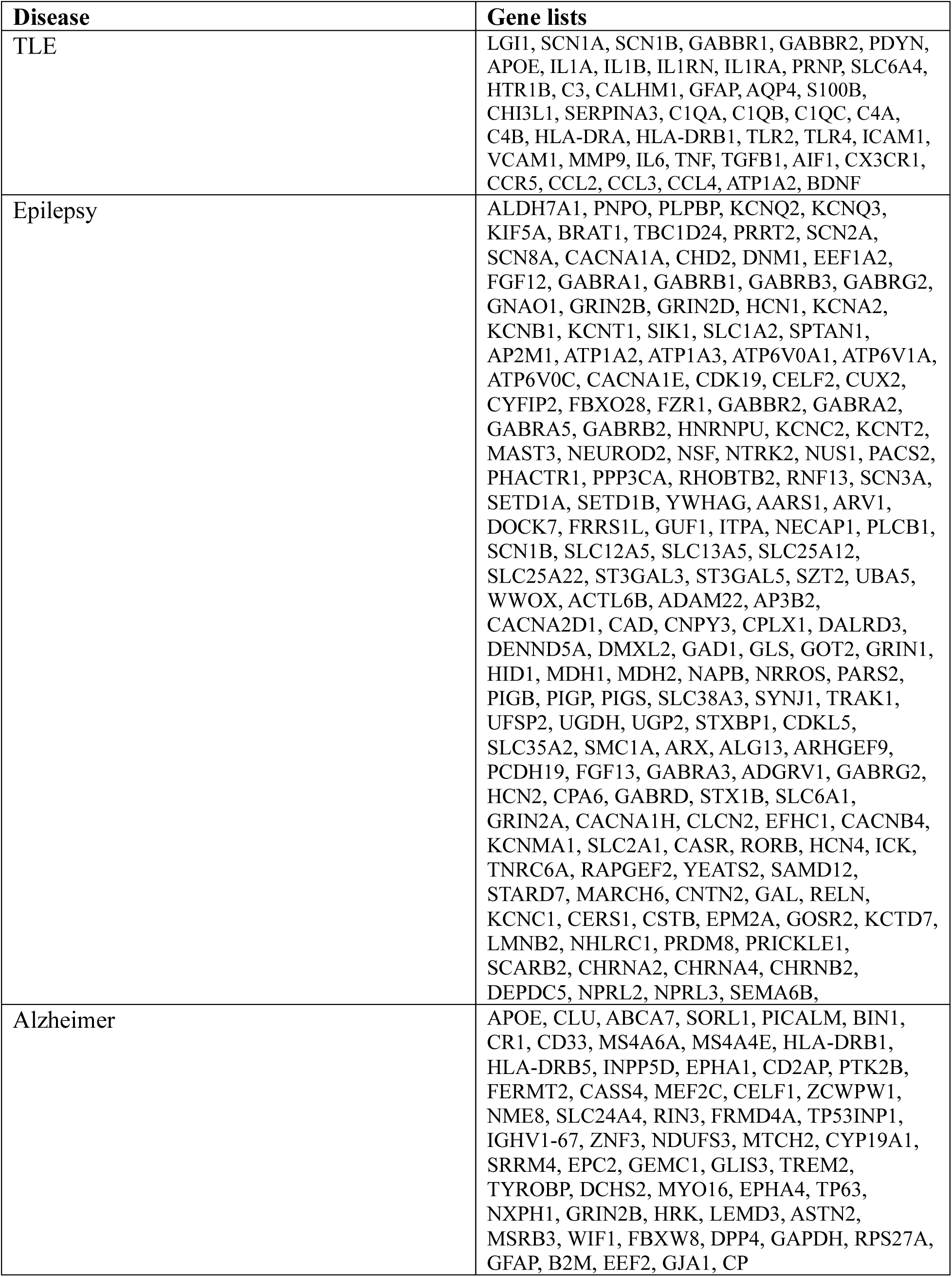
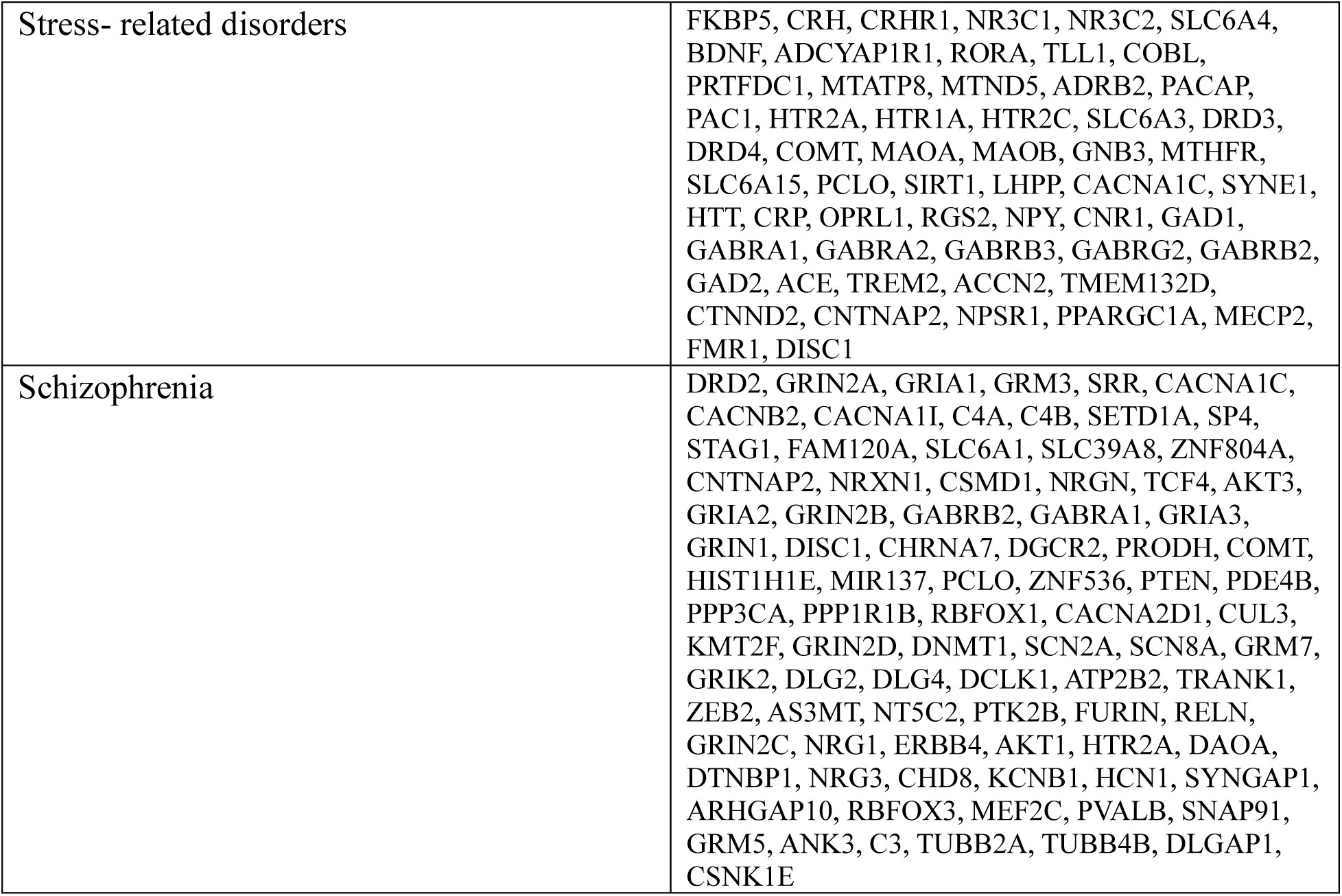
The lists of candidate genes used for disease specificity analysis.

